# Compromised beta-cell identity in type 2 diabetes

**DOI:** 10.1101/2023.03.20.533468

**Authors:** Pritha Dutta, Nadège Merabet, Rick Quax, Françoise Carlotti, Peter M.A. Sloot

## Abstract

Compromised beta-cell identity is emerging as an important contributor of beta-cell dysfunction in type 2 diabetes (T2D). Several studies suggest that hyperglycemia induces the inactivation of transcription factors involved in mature beta-cell identity. More specifically, chronic hyperglycemia leads to the downregulation of PDX1 and MAFA, two important beta-cell identity transcription factors and regulators of insulin promoter activity. Regulation of these transcription factors depends on interactions between multiple signaling cascades and processes driven by complex non-linear dynamics and taking place in different cellular compartments. To better understand these non-linear dynamics, we developed an integrated mathematical model of the underlying mechanisms regulating these key transcription factors. Our model was able to reproduce experimentally measured variations in the levels of PDX1, MAFA and insulin mRNA under different glucose concentrations. We used this model to simulate scenarios that could allow to restore PDX1 and MAFA levels and therefore insulin gene expression. From these simulations, FOXO1 emerged as an important target for the restoration of beta-cell identity.

**Author summary:** Glucose regulation depends on the secretion of insulin by beta-cells and uptake of glucose by the peripheral cells mediated by the action of insulin. In type 2 diabetes both aspects can be compromised. Defective insulin secretion results from compromised beta-cell function. One of the reasons behind compromised beta-cell function is that beta-cells fail to express one or more of the genes involved in insulin production and secretion and thus maintenance of beta-cell identity. The processes involved in the regulation of insulin production and secretion are complex. In this work, we are particularly interested in the role and downregulation of transcription factors, PDX1 and MAFA, which are critical regulators of insulin production, in relation with compromised beta-cell identity and function in the presence of chronic hyperglycemia. To understand better these complex processes, we use mathematical modelling which enables to generate hypotheses and simulate scenarios to extend our understanding of the mechanisms leading to compromised beta-cell function in the presence of chronic hyperglycemia. Our model and similar models can serve to identify therapeutical targets in beta-cells in order to restore their function.

## Introduction

Type 2 diabetes (T2D) is a chronic metabolic disorder characterized by defects in both insulin secretion and insulin action, which lead to supraphysiological plasma glucose levels. The beta-cells of the pancreatic islets play an important role in maintaining glucose homeostasis by secreting the hormone insulin, which contributes to lowering plasma glucose levels by promoting glucose uptake by skeletal muscle, liver and adipose tissue. Progressive deterioration in beta-cell function and decreased beta-cell mass have been observed in diabetic human islets. Post-mortem histological analysis of beta-cell mass of people with T2D has shown up to 50% reduction in beta-cell mass compared with healthy individuals with the same BMI [1–3]. Previously, this loss in beta-cell mass was entirely attributed to beta-cell apoptosis. However, multiple studies, both in animal models [4, 5] and in histological analyses of human pancreatic tissues [6–9], have demonstrated that compromised beta-cell identity might be an important contributor to the loss in functional beta-cell mass.

Maintenance of beta-cell identity requires active regulation of specific gene expression. Postmortem studies of pancreatic islets from diabetic patients have demonstrated a marked and selective loss of transcription factors involved in mature beta-cell identity including PDX1 and MAFA [6, 8, 9]. Since PDX1 and MAFA are also critical regulators of both insulin promoter activity and beta-cell function [10–12], their downregulation in response to chronic hyperglycemia is accompanied by a reduction in insulin gene expression and reduced insulin secretion [6, 9]. Loss of transcription factors PDX1 and MAFA is caused by a multitude of factors of which a leading factor is glucotoxicity [13]. Glucotoxicity is defined in terms of the detrimental effects of hyperglycemia on the phenotype and function of beta-cells.

Along with other causes, it has been found that two main processes contribute to glucotoxicity: oxidative stress and endoplasmic reticulum (ER) stress. Oxidative stress reflects an imbalance between the production of reactive oxygen species (ROS) and their elimination by antioxidant defenses. Chronic hyperglycemia increases the formation of reactive oxygen species (ROS) that, in excess and over time, can lead to chronic oxidative stress. Beta-cells are particularly vulnerable to oxidative stress due to the low expression of antioxidant enzymes such as catalase and glutathione peroxidase [14, 15]. In addition, chronic hyperglycemia places a high demand on the ER, which plays a major role in the production of insulin by synthesizing polypeptides from mRNA and converting them into mature proteins. A high inward flux of polypeptide molecules into the ER can overwhelm the protein-folding machinery, leading to an imbalance and the accumulation of unfolded and misfolded proteins, which is toxic for the cell. This imbalance is known as ER stress.

Oxidative stress and ER stress induced by chronic hyperglycemia have been shown to impair processes involved in the regulation of the beta-cell identity transcription factors PDX1 and MAFA. These processes are complex and non-linear and hence difficult to understand by only using experimental methods. Computational models can be useful tools to gain a comprehensive understanding of these intricate interactions by replicating the behaviour of the system based on known properties of the system components.

Numerous computational models in the literature simulate specific processes involved in beta-cell function. For instance, Chay and Keizer [16], Keizer and Magnus [17–19] and Smolen and Keizer [20, 21] proposed models of electrical activity in the beta-cells. Magnus and Keizer [18, 19, 22] further built models which include glucose metabolism, Ca^2+^ handling in the cytoplasm and mitochondria, and electrical activity in the plasma and inner mitochondrial membranes of the beta-cell. Bertram et al. [23] and Chay et al. [24] included the ER as a second Ca^2+^ compartment in beta-cell models. The literature contains many other models of glucose transport, glucose metabolism, Ca^2+^ handling, electrical activity, and glucose-stimulated insulin secretion [25–39]. However, in order to understand the mechanisms leading to compromised beta-cell identity and beta-cell dysfunction in the presence of chronic hyperglycemia, it is important to integrate these models of the different processes to obtain a more complete picture of the regulation of beta-cell function. These processes are interlinked and a perturbation to any of these processes can affect another process. In addition, these different models do not include processes related to the transcriptional activity involved in insulin production.

Hence, in order to further expand the understanding of hyperglycemia-induced beta-cell dysfunction, we developed a novel computational model which integrates key processes of beta-cell function and we proposed new equations for the regulation of PDX1, MAFA, and insulin transcription by stress-activated kinases and insulin signaling pathway. Our model was able to reproduce experimentally measured variations in the levels of PDX1, MAFA and insulin mRNA under different glucose concentrations. We found that with increasing glucose concentration, the levels of PDX1, MAFA and insulin mRNA decreased; they became almost negligible at the high glucose concentration of 30 mM. Furthermore, we used this model to simulate specific scenarios by targeting proteins that could allow to restore PDX1 and MAFA levels and therefore insulin gene expression. From these simulations, FOXO1 emerged as an important target for restoration of beta-cell identity, and therefore function. By integrating key processes involved in the maintenance of beta-cell identity and function, this model enables us to generate hypotheses and simulate scenarios to extend our understanding of the mechanisms leading to compromised beta-cell identity and beta-cell dysfunction in the presence of chronic hyperglycemia as well as identify potential intervention targets for the restoration of beta-cell identity.

## Results

### Model

Our model describes steady-states and dynamics of essential processes involved in the regulation of beta-cell function and identity. The different processes included encompass glucose transport into the beta-cell, glucose metabolism, calcium dynamics, electrical activity, protein folding, unfolded protein response (UPR), reactive oxygen species (ROS) production, and regulation of transcription factors necessary to maintain beta-cell identity. A schematic diagram of the model is given in Fig. 1. We modelled these processes using 31 coupled ordinary differential equations and 9 constraints (given in Table 1 and Table 2 respectively). These various processes are distributed over four different compartments in our model: the cytosol, mitochondria, the endoplasmic reticulum (ER) and the nucleus.

**Figure 1:**
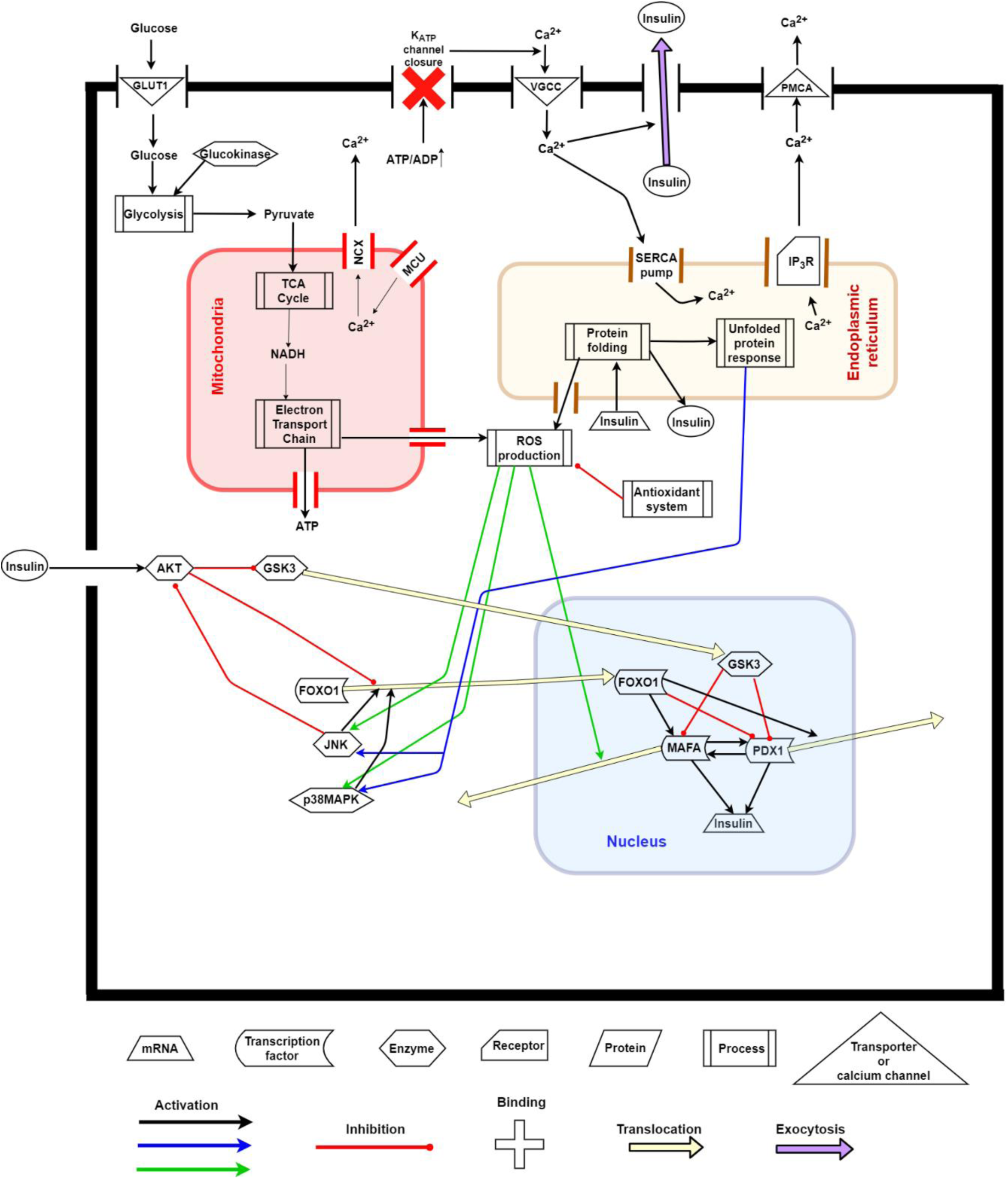
Schematic representation of the beta-cell model. The model includes four compartments: cytosol, mitochondria, endoplasmic reticulum (ER) and nucleus. Glucose enters the beta-cell through GLUT1 and then is metabolized through the processes of glycolysis, tricarboxylic acid (TCA) cycle and electron transport chain (ETC) which result in the production of ATP and reactive oxygen species (ROS). The rise in ATP/ADP ratio causes closure of the K-ATP channel and influx of Ca^2+^ into the beta-cell through the voltage-gated calcium channel (VGCC). Ca^2+^ leaves the beta-cell through the plasma membrane Ca^2+^ ATPase (PMCA). Ca^2+^ enters the mitochondria through the mitochondrial Ca^2+^ uniporter (MCU) complex and leaves through the mitochondrial Na^+^/Ca^2+^ exchanger (NCX). Ca^2+^ enters the ER through the sarco/endoplasmic reticulum Ca^2+^ ATPases (SERCA) and leaves through the inositol trisphosphate receptor (IP_3_R). Extra-cellular insulin activates AKT which in turn phosphorylates GSK3 and FOXO1 and prevents their translocation to the nucleus. On the other hand, ROS and the unfolded protein response (UPR) activate the stress kinases, JNK and p38MAPK, which in turn inhibit AKT and also promote the nuclear translocation of FOXO1. PDX1 and MAFA are the activator transcription factors of insulin as well as of each other. FOXO1 is a repressor and activator transcription factor of PDX1 and MAFA respectively. FOXO1 also causes nuclear exclusion of PDX1. GSK3 causes proteasomal degradation of both PDX1 and MAFA. Insulin mRNA then undergoes translation and enters the ER for folding. A high flux of insulin mRNA into the protein folding machinery activates the UPR. After folding, insulin is secreted from the beta-cell depending on the Ca^2+^ concentration in the cytosol.

**Table 1:**
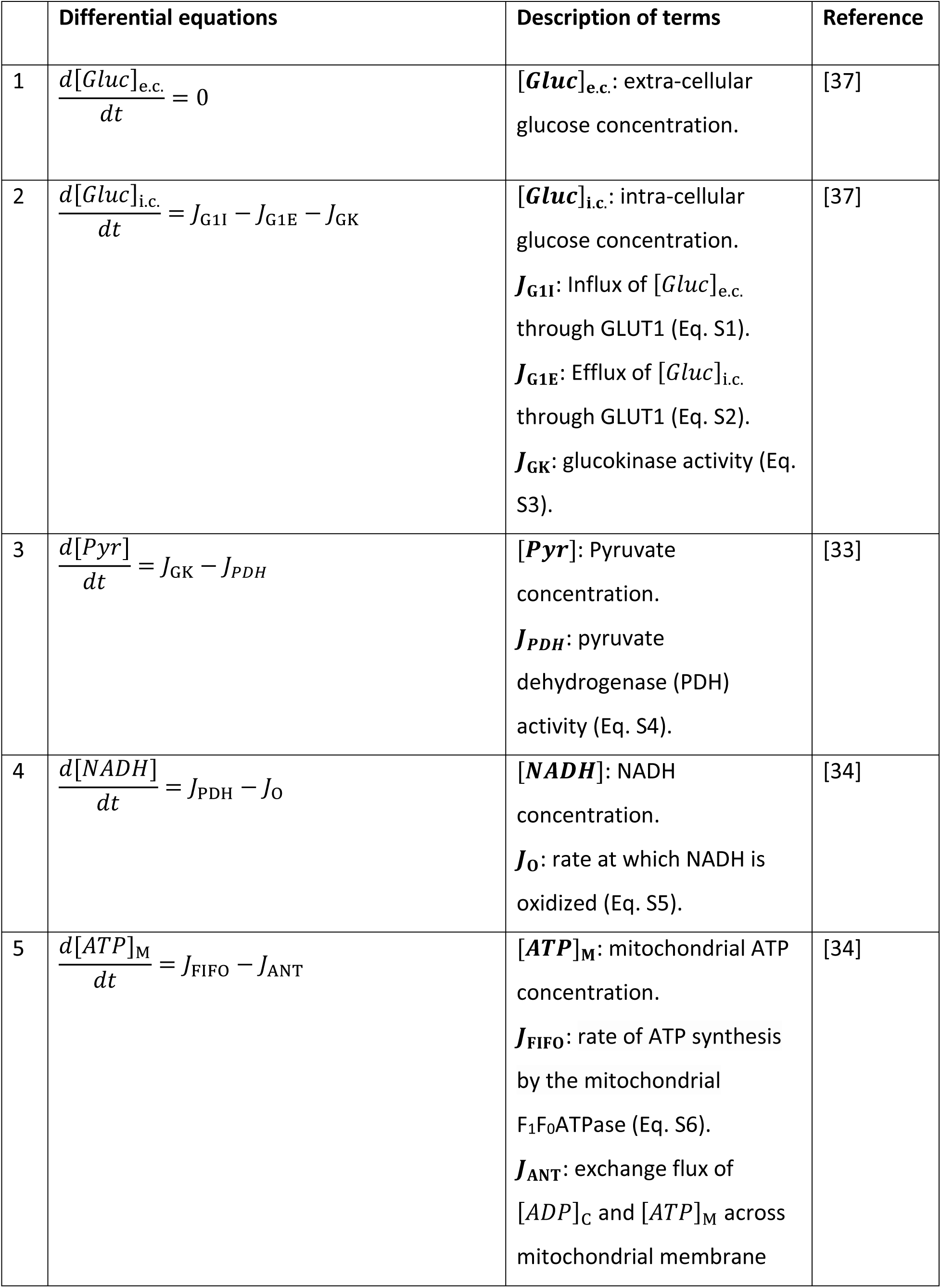
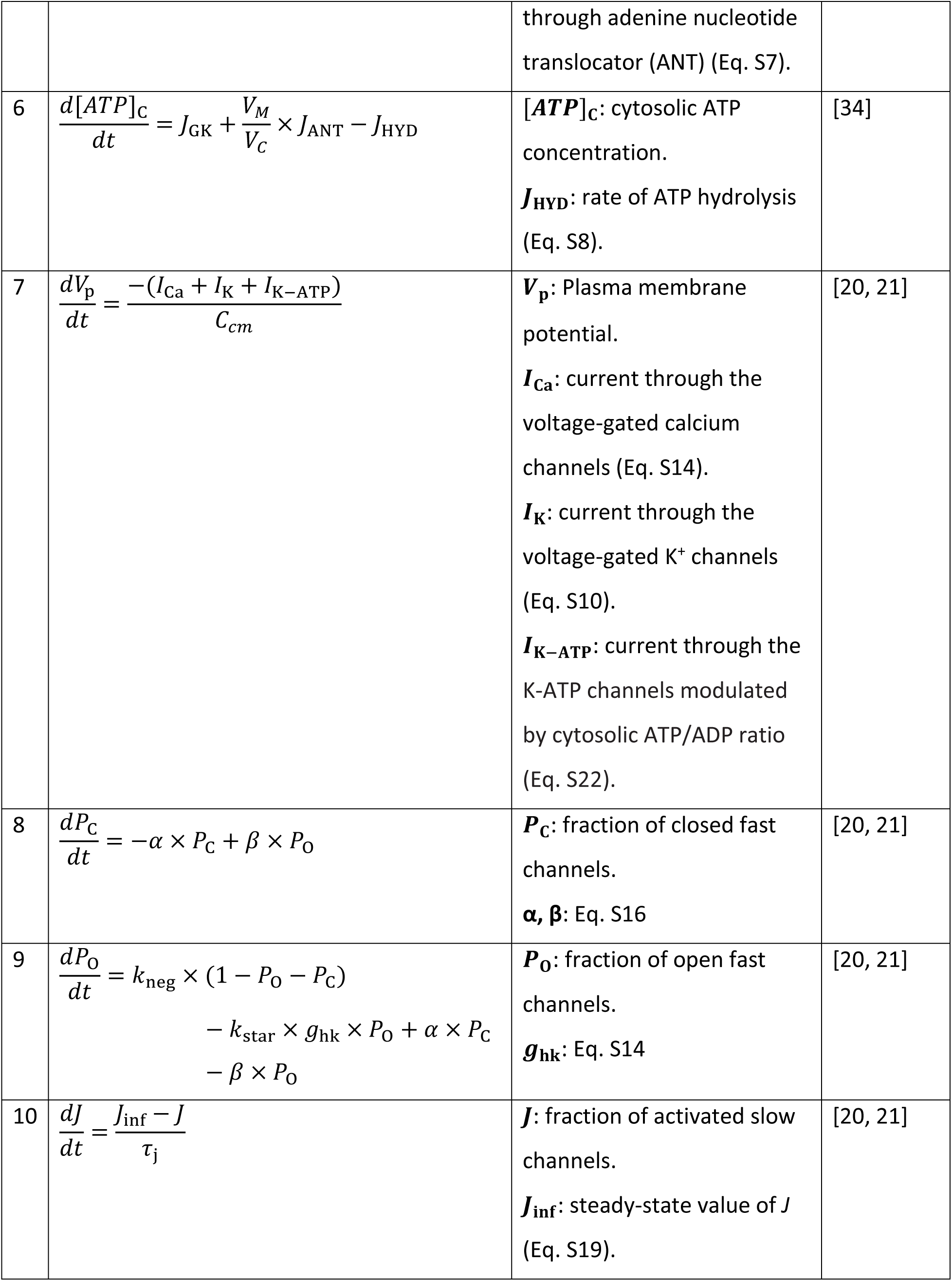
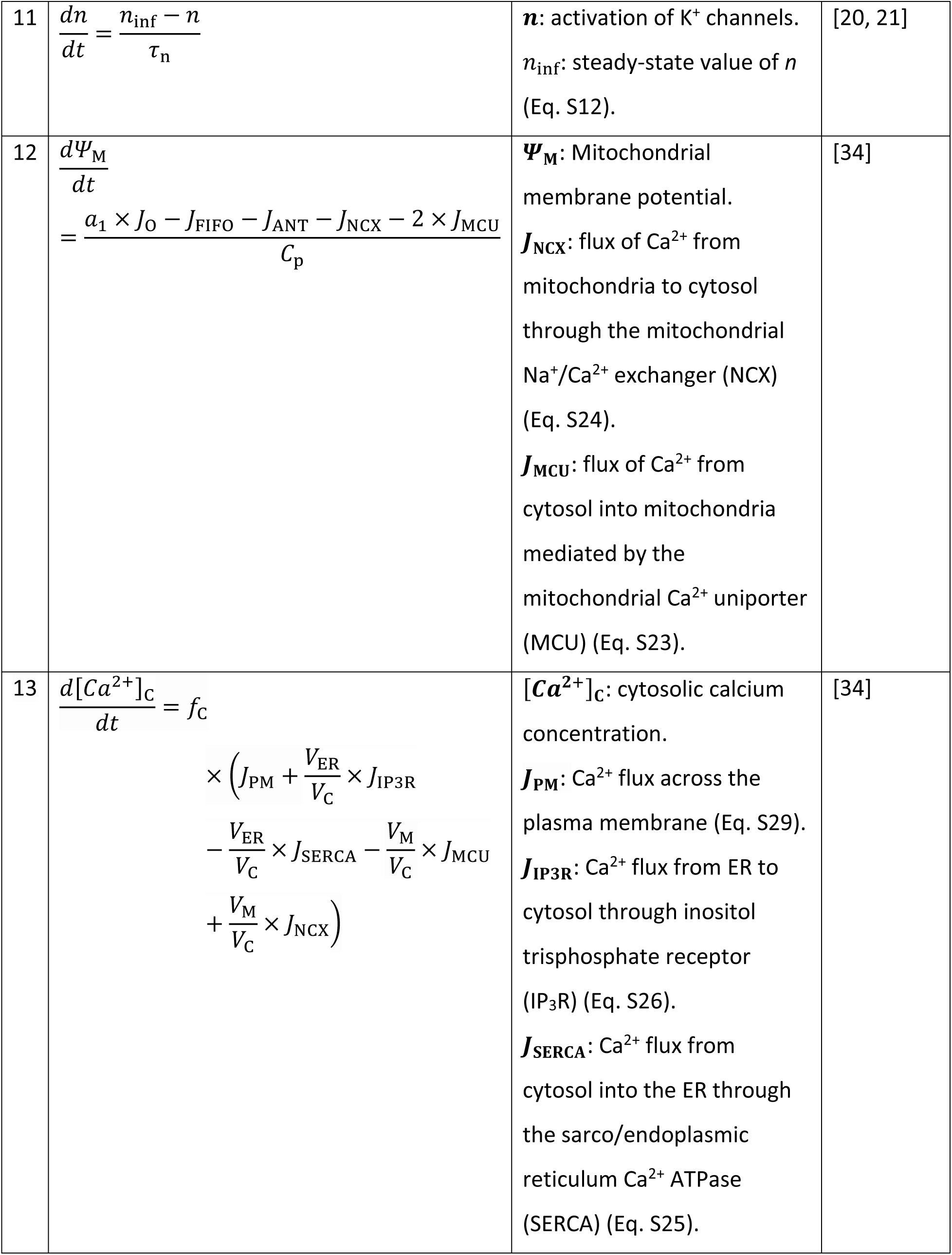
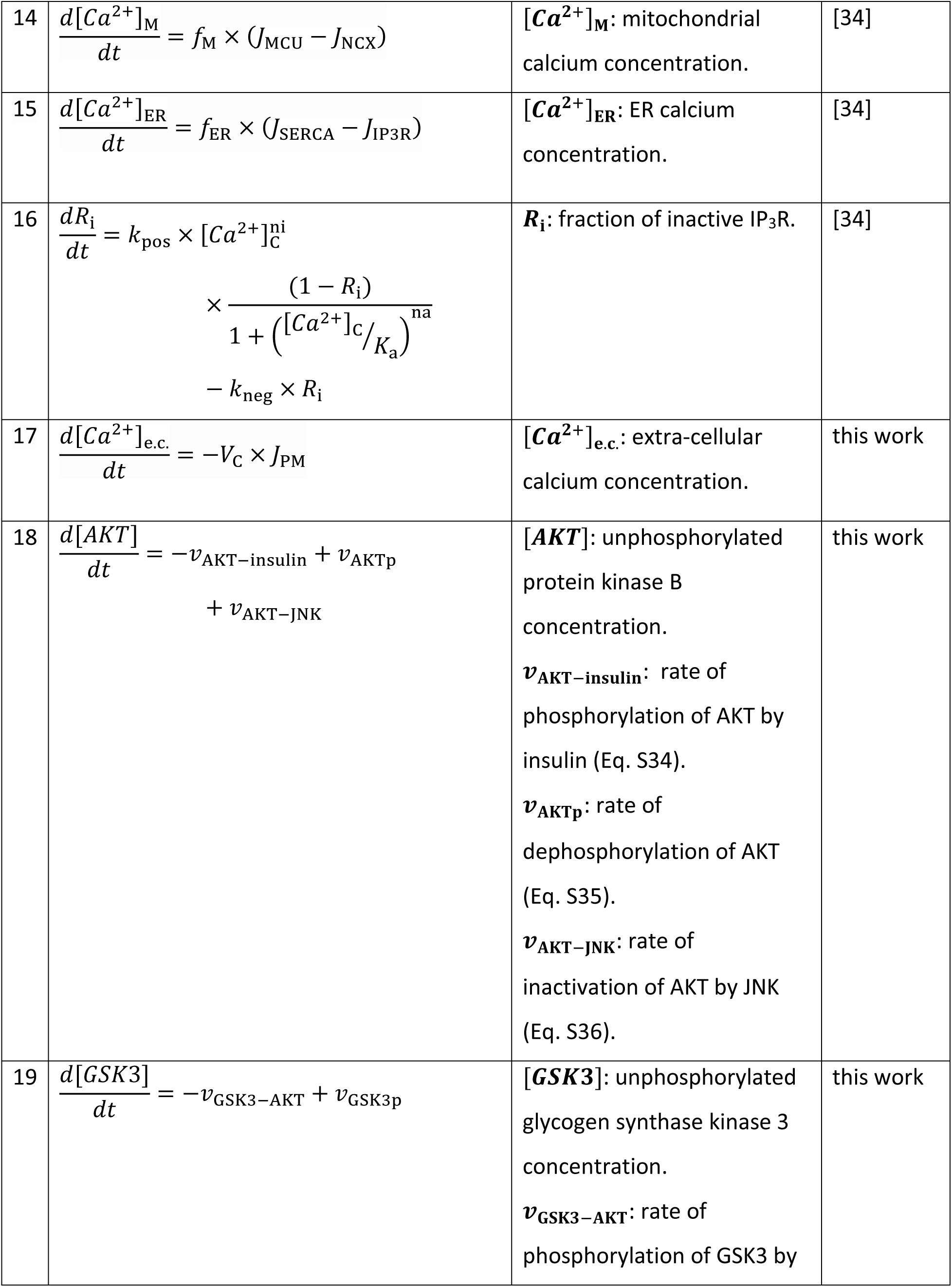
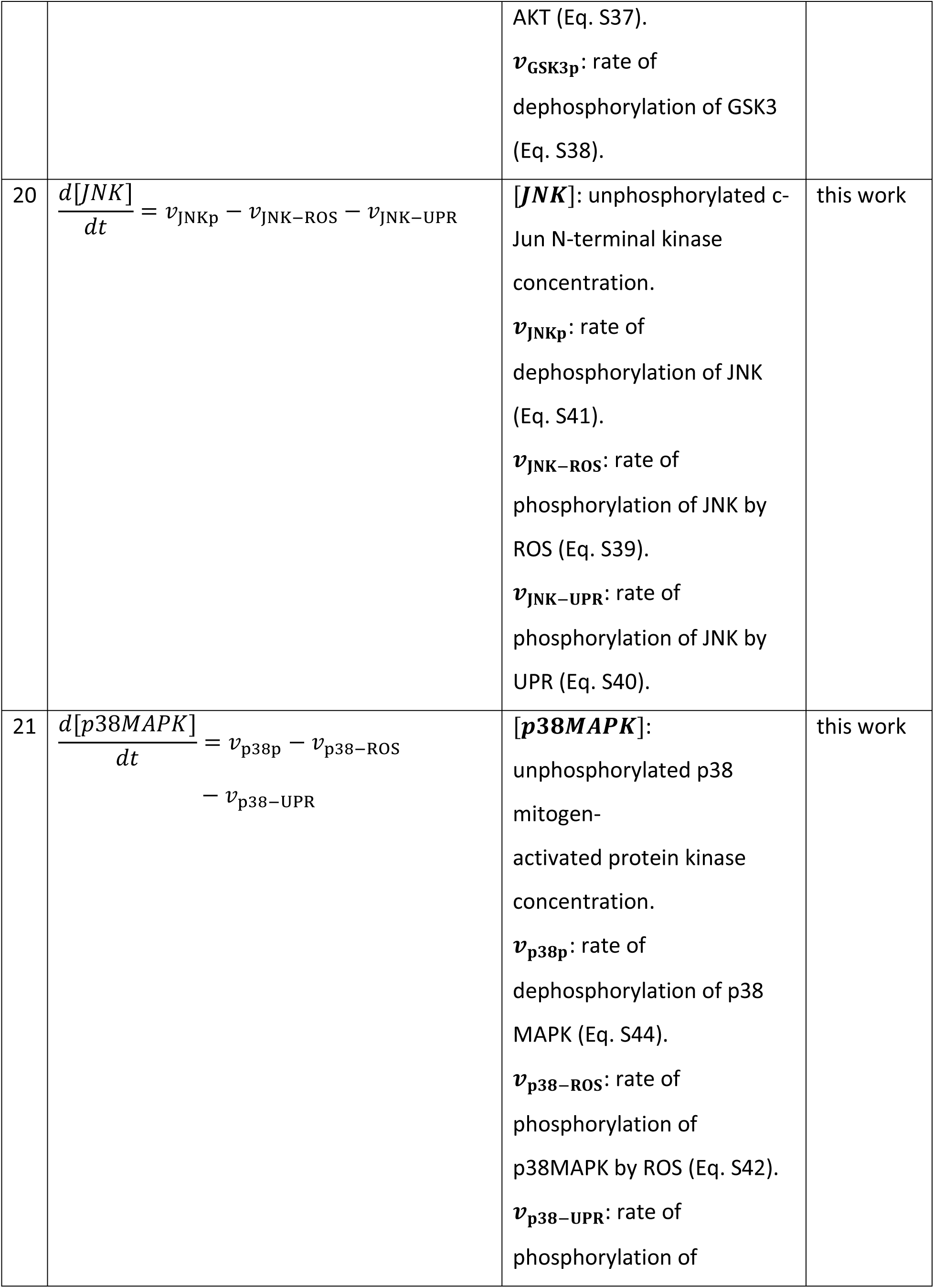
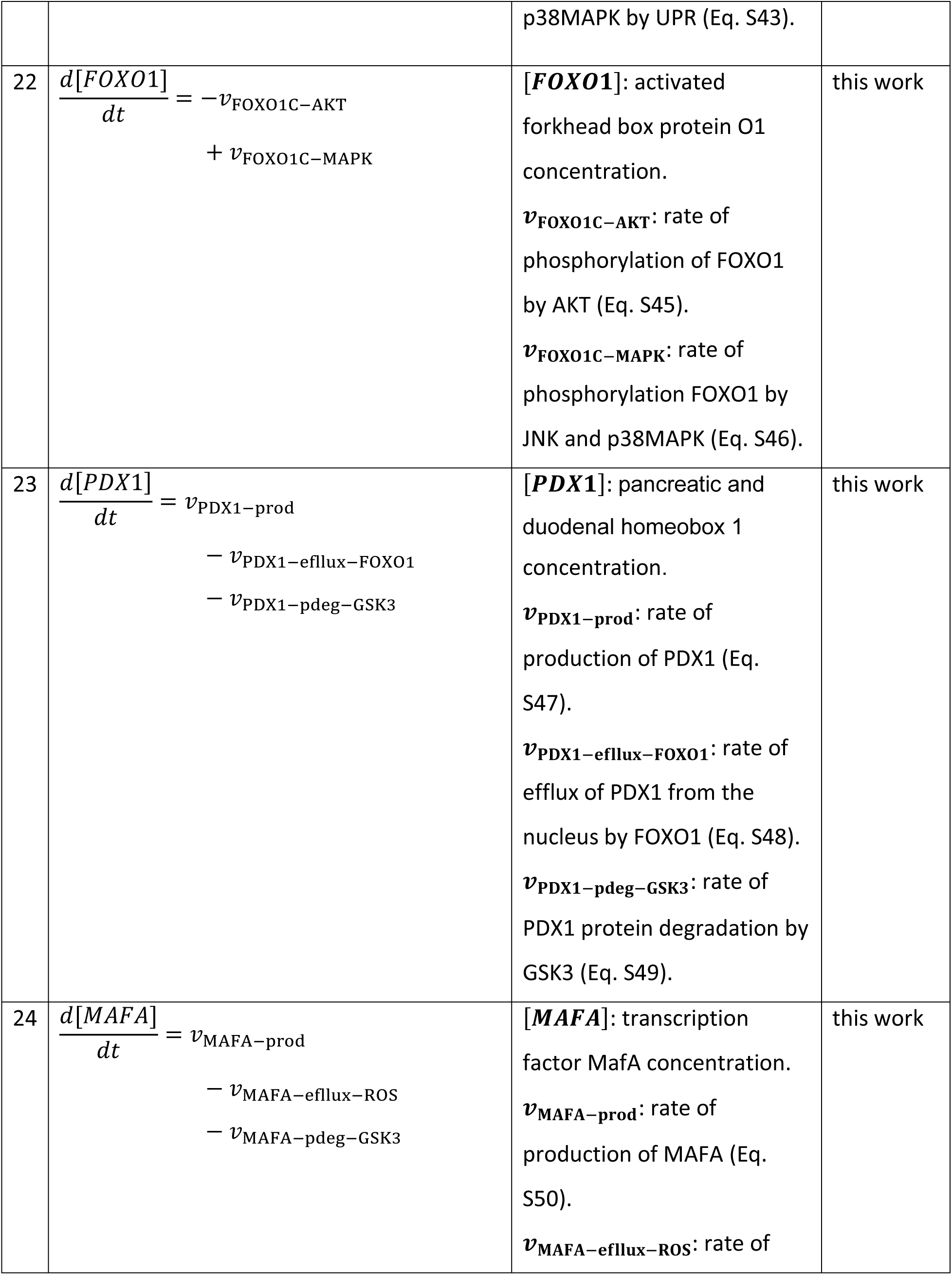
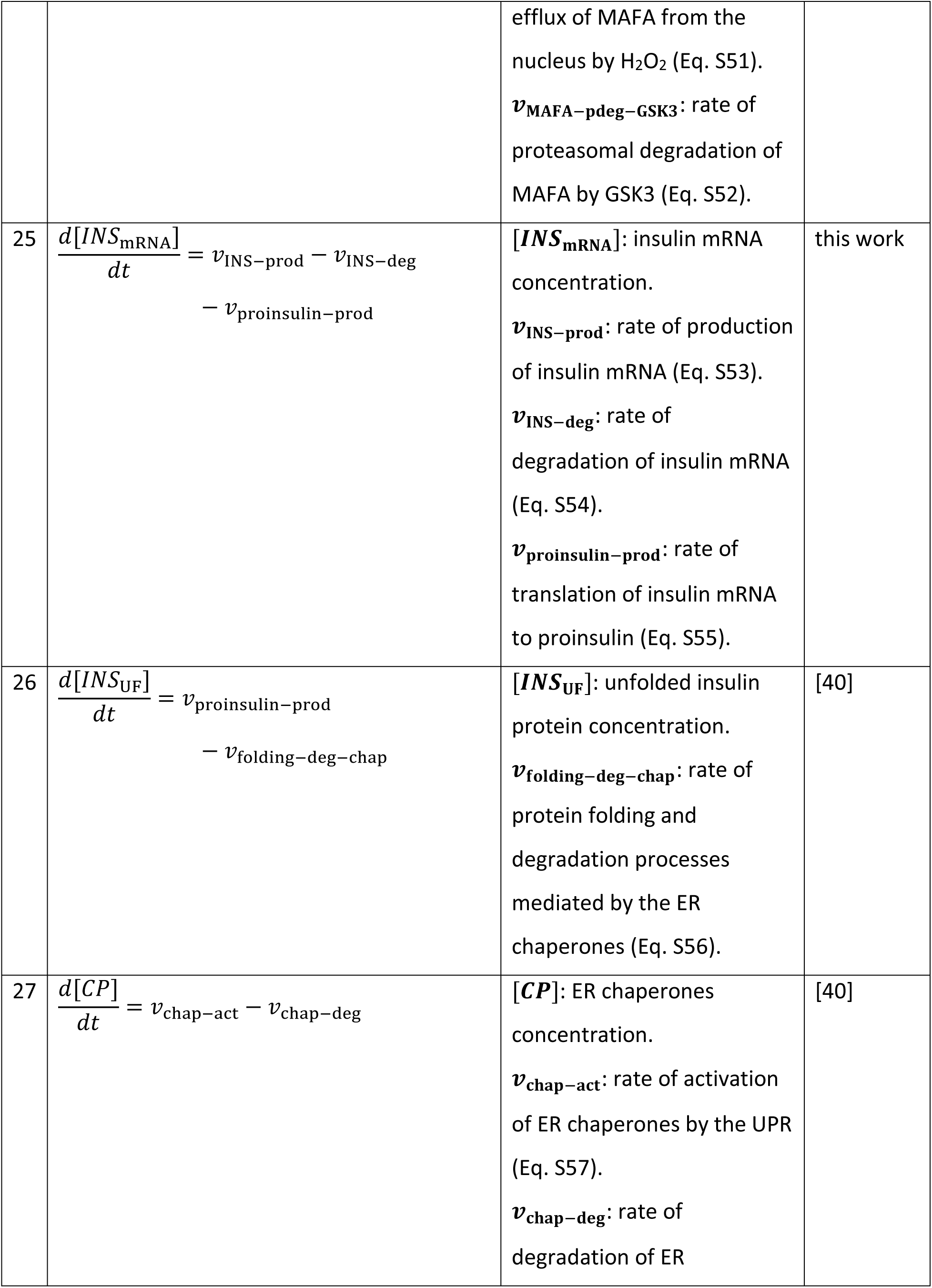
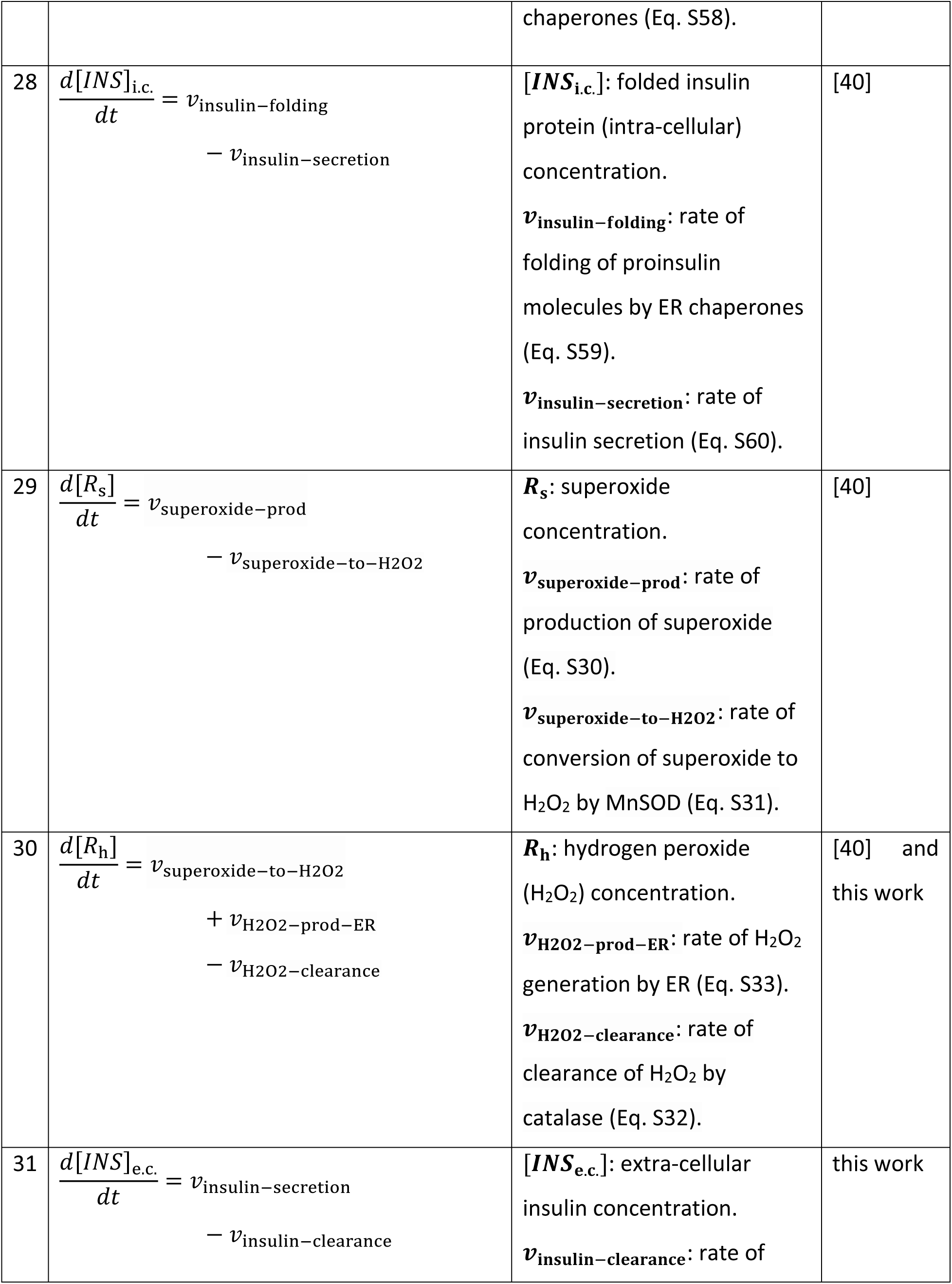
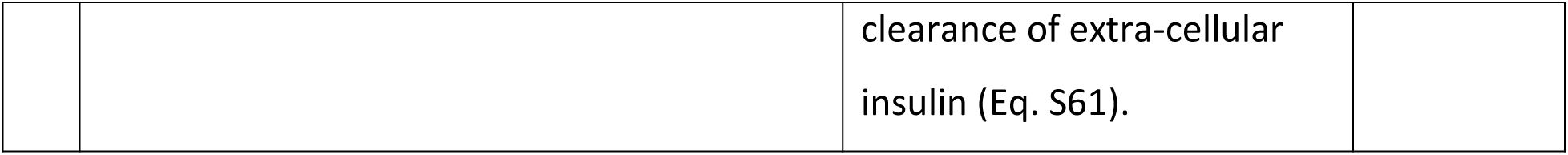
The system of differential equations used in the model. For details of all parameters used in the equations refer to Tables 3, 5 and 6. Equation numbers starting with S (eg. S1, S2) refer to equations in the Supporting Information file.

**Table 2:**
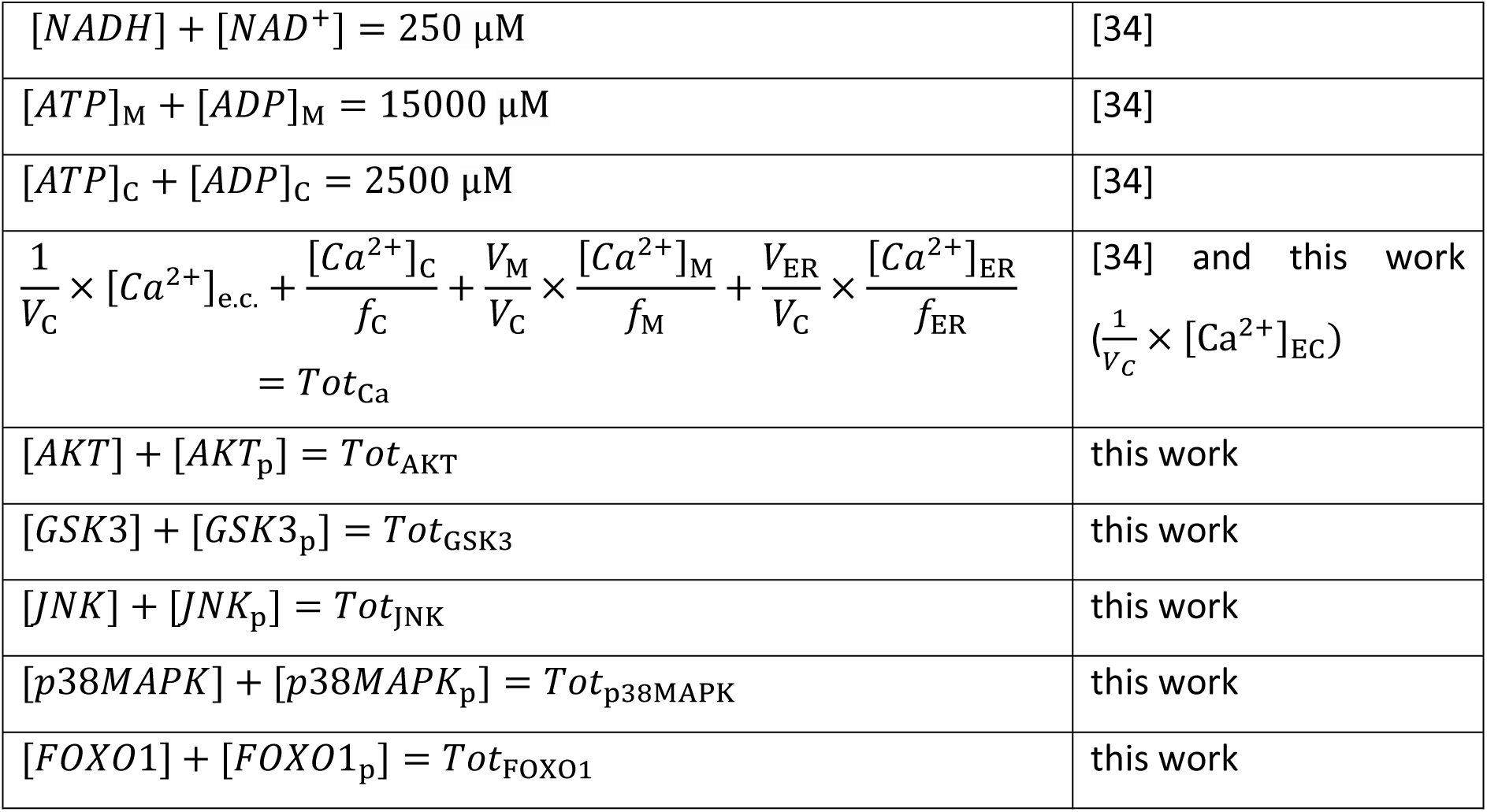
The constraints used in the model.

|  |  |
| --- | --- |
| $[NADH] + [NAD^+] = 250 \mu\text{M}$ | [34] |
| $[ATP]_M + [ADP]_M = 15000 \mu\text{M}$ | [34] |
| $[ATP]_C + [ADP]_C = 2500 \mu\text{M}$ | [34] |
| $\frac{1}{V_C} \times [Ca^{2+}]_{e.c.} + \frac{[Ca^{2+}]_C}{f_C} + \frac{V_M}{V_C} \times \frac{[Ca^{2+}]_M}{f_M} + \frac{V_{ER}}{V_C} \times \frac{[Ca^{2+}]_{ER}}{f_{ER}} = Tot_{Ca}$ | [34] and this work<br>($\frac{1}{V_C} \times [Ca^{2+}]_{EC}$) |
| $[AKT] + [AKT_p] = Tot_{AKT}$ | this work |
| $[GSK3] + [GSK3_p] = Tot_{GSK3}$ | this work |
| $[JNK] + [JNK_p] = Tot_{JNK}$ | this work |
| $[p38MAPK] + [p38MAPK_p] = Tot_{p38MAPK}$ | this work |
| $[FOXO1] + [FOXO1_p] = Tot_{FOXO1}$ | this work |

The rate equations describing the processes of glucose transport, glucose metabolism, calcium dynamics, electrical activity and protein folding are derived from the previous models of Luni et al. [37], Fridlyand et al. [33], Wacquier et al. [34], Chay et al. [16], Smolen et al. [20], and Graham [40]. We have added a term to the equation describing the rate of ATP hydrolysis (equation S8 in the **Supporting Information** file) from (Wacquier et al., 2016). The first term 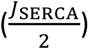 of the equation corresponds to the hydrolysis of ATP required to provide energy to the ER SERCA pumps to transport Ca^2+^ into the ER (Wacquier et al., 2016) while the added term (*k*_HYD_ × [*ATP*]_C_) represents all other ATP-consuming processes in the cytosol. We have also developed new equations to describe the regulation of PDX1, MAFA, and insulin transcription by stress-activated kinases and insulin signaling pathway (equations 18-25 in Table 1 and equations S34-S54 in the **Supporting Information** file). A detailed description of these different processes involved in beta-cell function and identity together with the rate equations used to represent these processes are provided in the **Supporting Information: Model development**.

### Parameter estimation

The values of the parameters used in the rate equations are extracted from the literature as well as estimated from experimental data. The parameters and the initial values of the state variables extracted from the literature are given in Tables 3 and 4 respectively.

**Table 3:** Model parameters and their values taken from the literature.

| Parameter | Description | Value | Reference |
| --- | --- | --- | --- |
| $V_C$ | Relative volume of cytosol w.r.t whole beta-cell volume | 0.882 | [41] |
| $V_N$ | Relative volume of nucleus w.r.t whole beta-cell volume | 0.117 | [41] |
| $V_{ER}$ | Relative volume of ER w.r.t whole beta-cell volume | 0.195 | [41] |
| $V_M$ | Relative volume of mitochondria w.r.t whole beta-cell volume | 0.039 | [41] |
| $K_{mG1}$ | Glucose concentration at which the reaction rate is half of $V_{mG1}$ | 3000 $\mu\text{M}$ | [37] |
| $K_{mATP}$ | Michaelis-Menten constant for the consumption of ATP in the glycolysis process | 500 $\mu\text{M}$ | [42] |
| $K_{GK}$ | Glucose concentration at which the reaction rate is half of $V_{mGK}$ | 7000 $\mu\text{M}$ | [42] |
| $n_{GK}$ | Hill coefficient | 1.7 | [42] |
| $q_1$ | Michaelis-Menten-like constant for $\text{NAD}^+$ consumption by the Krebs cycle | 1 | [34] |
| $q_2$ | $S_{0.5}$ value for activation the Krebs cycle by $\text{Ca}^{2+}$ | 0.1 $\mu\text{M}$ | [34] |
| $q_3$ | Michaelis-Menten constant for NADH consumption by the ETC | 100 $\mu\text{M}$ | [34] |
| $q_4$ | Voltage dependence coefficient 1 of ETC activity | 143 mV | [34] |
| $q_5$ | Voltage dependence coefficient 2 of ETC activity | 5 mV | [34] |
| $V_{F1FO}$ | Rate constant of the F1FO ATPase | 35000 $\mu\text{M.s}^{-1}$ | [34] |
| $q_6$ | Inhibition constant of ATPase activity by ATP | 10000 $\mu\text{M}$ | [34] |
| $q_7$ | Voltage dependence coefficient of ATPase activity | 150 mV | [34] |
| $q_8$ | Voltage dependence coefficient of ATPase activity | 8.5 mV | [34] |
| $V_{ANT}$ | Rate constant of the adenine nucleotide translocator | $5000 \mu\text{M.s}^{-1}$ | [34] |
| $\alpha_C$ | Factor taking cytosolic ADP and ATP buffering into account | 0.111 | [34] |
| $\alpha_M$ | Factor taking mitochondrial ADP and ATP buffering into account | 0.139 | [34] |
| $F'$ | $F/RT$; F: Faraday constant ($96480 \text{ C.mol}^{-1}$), R: Perfect gas constant ($8315 \text{ mJ.mol}^{-1}.\text{K}^{-1}$), T: Temperature (310.16 K) | $0.037 \text{ mV}^{-1}$ | [34] |
| $V_{IP3R}$ | Maximum $\text{Ca}^{2+}$ flux through $\text{IP}_3\text{R}$ | $0.1986 \text{ s}^{-1}$ | [34] |
| $K_{IP3}$ | Dissociation constant of $\text{IP}_3$ binding from its receptor | $1 \mu\text{M}$ | [34] |
| $K_a$ | Dissociation constant of $\text{Ca}^{2+}$ from the activating site of the $\text{IP}_3\text{R}$ | $0.3 \mu\text{M}$ | [34] |
| $n_a$ | Hill coefficient of $\text{Ca}^{2+}$ binding to the activating site of the $\text{IP}_3\text{R}$ | 3 | [34] |
| $V_{SERCA}$ | Maximum $\text{Ca}^{2+}$ flux through SERCA | $120 \mu\text{M.s}^{-1}$ | [34] |
| $K_p$ | Dissociation constant of $\text{Ca}^{2+}$ from SERCA | $0.35 \mu\text{M}$ | [34] |
| $K_e$ | Dissociation constant of ATP from SERCA pumps | $0.05 \mu\text{M}$ | [34] |
| $V_{MCU}$ | Maximum $\text{Ca}^{2+}$ flux through MCU | $0.0006 \mu\text{M.s}^{-1}$ | [34] |
| $K_{MCU}$ | Half-maximal activation constant for MCU | $0.9682 \mu\text{M}$ | [34] |
| $p_1$ | Voltage dependence coefficient of MCU activity | $0.1 \text{ mV}^{-1}$ | [34] |
| $V_{NCX}$ | Maximum $\text{Ca}^{2+}$ flux through NCX | $0.35 \mu\text{M.s}^{-1}$ | [34] |
| $p_2$ | Voltage dependence coefficient of NCX activity | $0.016 \text{ mV}^{-1}$ | [34] |
| $C_p$ | Mitochondrial inner membrane capacitance divided by F | $1.8 \mu\text{M.mV}^{-1}$ | [34] |
| $f_C$ | Calcium buffering factor in cytosol | 0.01 | [34] |
| $f_M$ | Calcium buffering factor in mitochondria | 0.0003 | [34] |
| $f_{ER}$ | Calcium buffering factor in endoplasmic reticulum | 0.01 | [34] |
| $k_{\text{pos}}$ | Rate constant of $\text{Ca}^{2+}$ binding to the inhibiting site of the $\text{IP}_3\text{R}$ | $20 \mu\text{M}^{-4}.\text{s}^{-1}$ | [34] |
| $k_{\text{neg}}$ | Rate constant of $\text{Ca}^{2+}$ dissociation from the inactivating site of the $\text{IP}_3\text{R}$ | $0.02 \text{ s}^{-1}$ | [34] |
| $n_i$ | Hill coefficient of $\text{Ca}^{2+}$ binding to the inhibiting site of the $\text{IP}_3\text{R}$ | 4 | [34] |
| $C_{\text{pm}}$ | Cell membrane capacitance | $5310 (\mu\text{F}/\text{cm}^2)$ | [16] |
| $k_{\text{dmINS}}$ | Rate of insulin mRNA degradation | $0.000006637 \text{ s}^{-1}$ | [43] |
| $k_{\text{dmINS}}$ | Rate of translation of insulin mRNA | $0.035584 \text{ s}^{-1}$ | [40] |
| $d_{\text{uu}}$ | Rate of proinsulin folding/degradation | $0.067 \text{ s}^{-1}$ | [40] |
| $K_{\text{uu}}$ | Proinsulin-chaperones Michaelis-Menten constant (folding/degradation) | $5.93 \times 10^{-2} \mu\text{M}$ | [40] |
| $d_{\text{CP}}$ | Rate of chaperones decay | $0.2918 \text{ s}^{-1}$ | [40] |
| $q_f$ | Proinsulin folding fraction | 0.8 | [40] |
| $k_{\text{IS}}$ | Rate of insulin secretion | $0.0002041 \text{ s}^{-1}$ | [40] |
| $k_{\text{IC}}$ | Rate of insulin clearance | $351.115 \text{ s}^{-1}$ | [40] |
| $K_1$ | PERK-proinsulin dissociation constant | 5.93 $\mu\text{M}$ | [40] |
| $K_2$ | PERK-chaperones dissociation constant | $3.56 \times 10^{-2} \mu\text{M}$ | [40] |
| $K_3$ | Proinsulin-chaperones dissociation constant | 1.19 $\mu\text{M}$ | [40] |
| $q_r$ | Fractional superoxide production | 9.76612 | [40] |
| $k_{rs}$ | Superoxide dismutation rate | $2400 \mu\text{M}^{-1}\text{s}^{-1}$ | [40] |
| $A_s$ | MnSOD concentration | 5 $\mu\text{M}$ | [40] |
| $\delta_{rh}$ | Rate of hydrogen peroxide removal by catalase | $17 \mu\text{M}^{-1}\text{s}^{-1}$ | [40] |
| $A_h$ | Catalase concentration | 3 $\mu\text{M}$ | [40] |

**Table 4:** Initial values of model state variables taken from the literature.

| Variable | Description | Initial value | Reference |
| --- | --- | --- | --- |
| $[Pyr]$ | Pyruvate | 9 $\mu\text{M}$ | [33] |
| $[NADH]$ | NADH | 60 $\mu\text{M}$ | [44] |
| $[ATP]_C$ | Cytosolic ATP | 1400 $\mu\text{M}$ | [34] |
| $[ATP]_M$ | Mitochondrial ATP | 11250 $\mu\text{M}$ | [34] |
| $\Psi_M$ | Mitochondrial membrane potential | 130 mV | [45] |
| $[Ca^{2+}]_C$ | Cytosolic calcium | 0.1 $\mu\text{M}$ | [46, 47] |
| $[Ca^{2+}]_M$ | Mitochondrial calcium | 0.1 $\mu\text{M}$ | [47, 48] |
| $[Ca^{2+}]_{ER}$ | ER calcium | 200 $\mu\text{M}$ | [49, 50] |
| $[Ca^{2+}]_{e.c.}$ | Extra-cellular calcium | 2000 $\mu\text{M}$ | [51] |
| $R_i$ | Fraction of inactive $IP_3$ receptors | 0.088 | [34] |
| $V_p$ | Plasma membrane potential | -64 mV | [20] |
| $P_C$ | Fraction of closed fast channels | 1 | [20] |
| $P_O$ | Fraction of open fast channels | 0 | [20] |
| $J$ | Fraction of activated slow channels | 0.86 | [20] |
| $n$ | Fraction of activated $K^+$ channel | 0 | [20] |
| $[INS_{mRNA}]$ | Insulin mRNA concentration | 1.3424 $\mu M$ | [40] |
| $[INS_{UF}]$ | Unfolded insulin protein concentration | 5.93 $\mu M$ | [40] |
| $[Chap]$ | ER chaperones | 0.593 $\mu M$ | [40] |
| $[INS]_{i.c.}$ | Folded insulin protein concentration | 189.234 $\mu M$ | [40] |
| $[R_s]$ | Superoxide | 0.01 $\mu M$ | [40] |
| $[R_h]$ | Hydrogen peroxide ($H_2O_2$) | 0.1 $\mu M$ | [40] |
| $[INS]_{e.c.}$ | Extra-cellular insulin | 0.00009 $\mu M$ | [40] |

The parameter estimation was done in two stages. The first stage of the parameter estimation involved the processes of glucose transport, glucose metabolism, electrical activity, and calcium dynamics and corresponded to the estimation of 8 parameters. This part was parameterized separately as a first step because it does not have any feedback from the rest of the model. These estimated parameters from the first stage were then used as inputs to the second stage of the parameter estimation which included the whole model. The number of unknown parameters for the second stage of estimation was 37.

The first stage of the parameter estimation was repeated 100 times. The estimated parameters are presented in Table 5 as mean ± standard deviation (SD) (n = 100). Fig. 2 shows the comparison of the simulated data with the experimental data. The details of the experimental data and the cost function equations are given in the **Methods** section.

**Figure 2:**
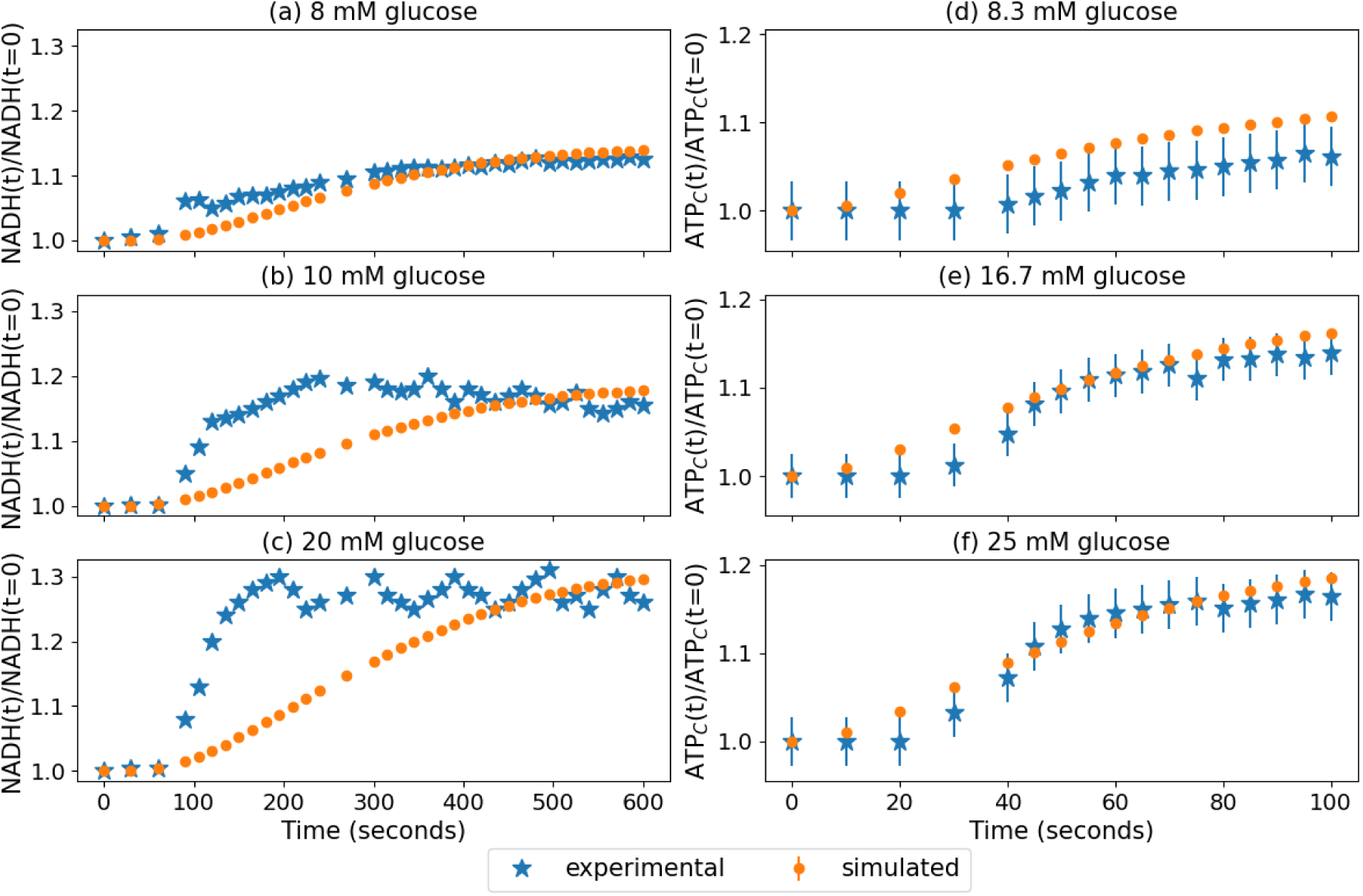
Stage 1 parameter estimation results: comparison of simulated data with experimental data. The simulated data are presented as mean ± SD (n = 100).

**Table 5:** Details of the estimated parameters from stage 1 of the parameter estimation. The values are presented as mean ± SD (n = 100).

| | Parameter | Parameter description | Mean ± SD |
| --- | --- | --- | --- |
| 1 | $V_{mG1}$ | Maximum reaction rate for GLUT1 that is observed at saturating glucose concentrations | $50.86 \pm 0.93 \mu\text{M} \cdot \text{s}^{-1}$ |
| 2 | $V_{mGK}$ | Maximum rate of glucose consumption | $375.55 \pm 15.51 \mu\text{M} \cdot \text{s}^{-1}$ |
| 3 | $V_{mPDH}$ | Maximum rate of PDH reaction | $30.36 \pm 0.15 \mu\text{M} \cdot \text{s}^{-1}$ |
| 4 | $V_O$ | Maximum rate of NADH oxidation by ETC | $32.88 \pm 0.38 \mu\text{M} \cdot \text{s}^{-1}$ |
| 5 | $K_{mPyr}$ | Michaelis-Menten constant for oxidation of pyruvate | $0.055 \pm 0.008 \mu\text{M}$ |
| 6 | $k_{Hyd}$ | Rate of ATP hydrolysis | $0.056 \pm 0.0005 \text{ s}^{-1}$ |
| 7 | $k_{PMCA}$ | Rate constant for the removal of $[\text{Ca}^{2+}]_c$ through plasma membrane $\text{Ca}^{2+}$ ATPase (PMCA) | $37.99 \pm 0.14 \text{ s}^{-1}$ |
| 8 | $a_1$ | Scaling factor between NADH consumption and change in membrane voltage | $249.96 \pm 0.1$ |

The second stage of the parameter estimation was repeated 100 times and the estimated parameters are presented in Table 6 as mean ± SD (n = 100). Fig. 3 shows the comparison of the simulated data with the experimental data. The details of the experimental data and the cost function equations are given in the **Methods** section.

**Figure 3:**
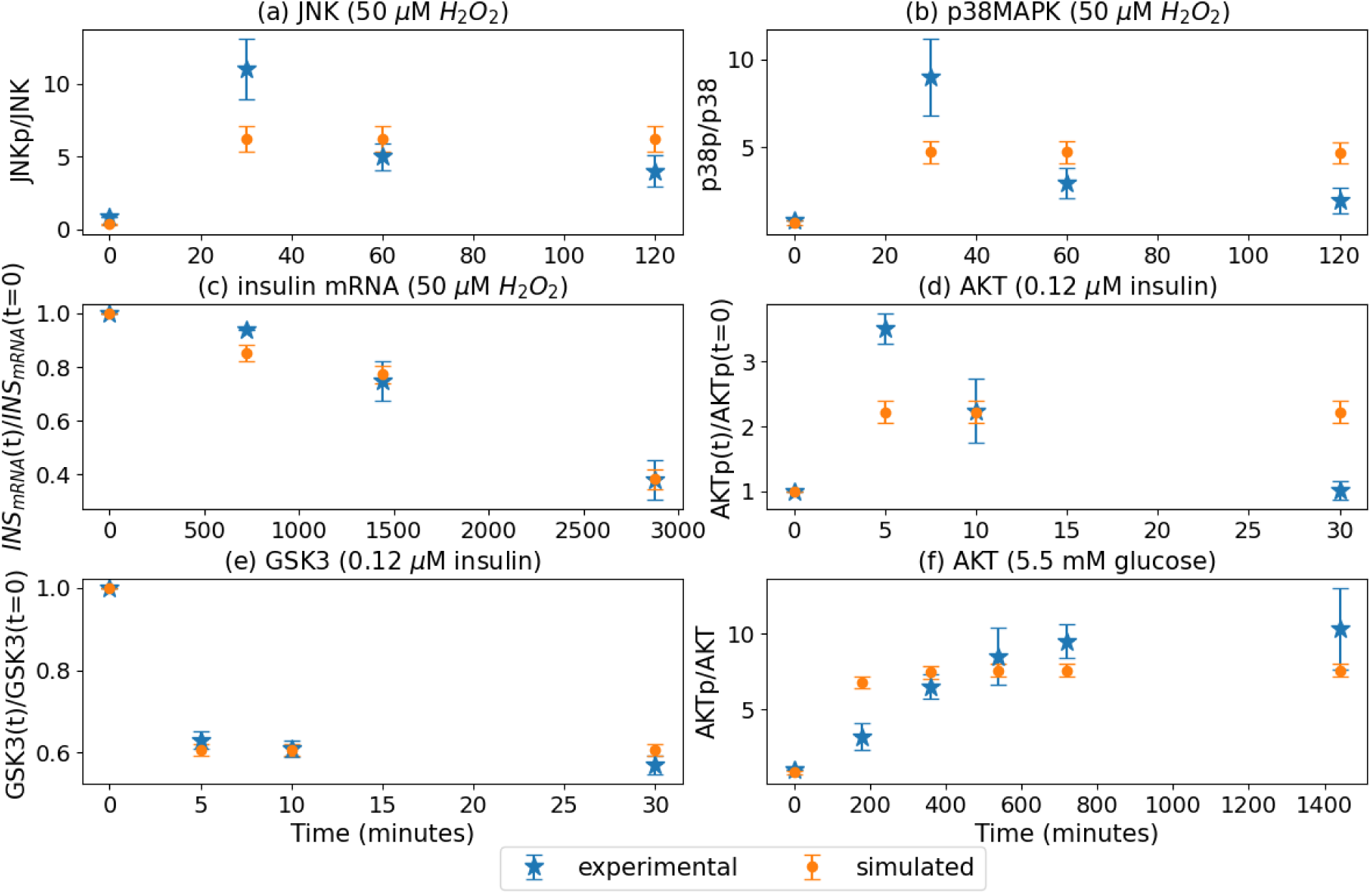
Stage 2 parameter estimation results: comparison of simulated data with experimental data. The simulated data is presented as mean ± SD (n = 100).

**Table 6:** Details of the estimated parameters from stage 2 of the parameter estimation. The values are presented as mean ± SD (n = 100).

| | Parameter | Parameter description | Mean ± SD |
| --- | --- | --- | --- |
| 1 | $[AKT]_{t=0}$ | Initial concentration of unphosphorylated AKT | $3.98 \pm 0.55 \mu\text{M}$ |
| 2 | $[AKT_p]_{t=0}$ | Initial concentration of phosphorylated AKT | $3.22 \pm 0.49 \mu\text{M}$ |
| 3 | $[GSK3]_{t=0}$ | Initial concentration of unphosphorylated GSK3 | $3.1 \pm 0.58 \mu\text{M}$ |
| 4 | $[GSK3_p]_{t=0}$ | Initial concentration of phosphorylated GSK3 | $3.45 \pm 0.52 \mu\text{M}$ |
| 5 | $[JNK]_{t=0}$ | Initial concentration of unphosphorylated JNK | $4.65 \pm 0.25 \mu\text{M}$ |
| 6 | $[JNK_p]_{t=0}$ | Initial concentration of phosphorylated JNK | $1.64 \pm 0.24 \mu\text{M}$ |
| 7 | $[p38MAPK]_{t=0}$ | Initial concentration of unphosphorylated p38MAPK | $4.62 \pm 0.28 \mu\text{M}$ |
| 8 | $[p38MAPK_p]_{t=0}$ | Initial concentration of phosphorylated p38MAPK | $3.19 \pm 0.57 \mu\text{M}$ |
| 9 | $[FOXO1]_{t=0}$ | Initial concentration of unphosphorylated FOXO1 | $3.69 \pm 0.22 \mu\text{M}$ |
| 10 | $[FOXO1_p]_{t=0}$ | Initial concentration of phosphorylated FOXO1 | $3.68 \pm 0.23 \mu\text{M}$ |
| 11 | $[PDX1]_{t=0}$ | Initial concentration of PDX1 | $3.27 \pm 0.5 \mu\text{M}$ |
| 12 | $[MAFA]_{t=0}$ | Initial concentration of MAFA | $3.61 \pm 0.27 \mu\text{M}$ |
| 13 | $k_{AKT}$ | Phosphorylation rate of AKT by insulin | $4.19 \pm 0.55 \mu\text{M}$ |
| 14 | $k_{AKT_p}$ | AKT dephosphorylation rate | $0.33 \pm 0.13 \mu\text{M}$ |
| 15 | $k_{AKT-JNK}$ | Rate of inactivation of AKT by JNK | $0.24 \pm 0.034 \mu\text{M}$ |
| 16 | $k_{GSK3}$ | Phosphorylation rate of GSK3 by AKT | $0.3 \pm 0.056 \mu\text{M}$ |
| 17 | $k_{GSK3_p}$ | GSK3 dephosphorylation rate | $0.85 \pm 0.11 \mu\text{M}$ |
| 18 | $k_{JNK-ROS}$ | Phosphorylation rate of JNK by $H_2O_2$ | $4.71 \pm 0.21 \mu\text{M}$ |
| 19 | $k_{JNK-UPR}$ | Phosphorylation rate of JNK by UPR | $4.17 \pm 0.76 \mu\text{M}$ |
| 20 | $k_{p38-ROS}$ | Phosphorylation rate of p38MAPK by $H_2O_2$ | $3.53 \pm 0.68 \mu\text{M}$ |
| 21 | $k_{p38-UPR}$ | Phosphorylation rate of p38MAPK by UPR | $5.2 \pm 0.65 \mu\text{M}$ |
| 22 | $k_{MAPK_p}$ | Dephosphorylation rate of JNK/p38MPK | $1.67 \pm 0.2 \mu\text{M}$ |
| 23 | $k_{\text{FOXO1-AKTp}}$ | Phosphorylation rate of FOXO1 by AKT | $1.66 \pm 0.2 \mu\text{M}$ |
| 24 | $k_{\text{FOXO1p-MAPKp}}$ | Phosphorylation rate of FOXO1 by JNK and p38MAPK | $1.14 \pm 0.23 \mu\text{M}$ |
| 25 | $V_{\text{mPDX1}}$ | Maximum rate of PDX1 mRNA production | $1.53 \pm 0.27 \mu\text{M}$ |
| 26 | $K_{\text{MAFA-PDX1}}$ | Dissociation constant of MAFA from the binding site of PDX1 promoter | $2.49 \pm 0.43 \mu\text{M}$ |
| 27 | $K_{\text{FOXO1-PDX1}}$ | Dissociation constant of FOXO1 from the binding site of PDX1 promoter | $3.67 \pm 0.85 \mu\text{M}$ |
| 28 | $k_{\text{PDX1-FOXO1-efflux}}$ | Rate of efflux of PDX1 from nucleus by FOXO1 | $0.14 \pm 0.024 \mu\text{M}$ |
| 29 | $k_{\text{GSK3-PDX1-pdeg}}$ | Rate of PDX1 protein degradation by GSK3 | $0.093 \pm 0.007 \mu\text{M}$ |
| 30 | $V_{\text{mMAFA}}$ | Maximum rate of MAFA mRNA production | $2.02 \pm 0.49 \mu\text{M}$ |
| 31 | $K_{\text{PDX1-MAFA}}$ | Dissociation constant of PDX1 from the binding site of MAFA promoter | $2.67 \pm 0.21 \mu\text{M}$ |
| 32 | $K_{\text{FOXO1-MAFA}}$ | Dissociation constant of FOXO1 from the binding site of MAFA promoter | $0.62 \pm 0.22 \mu\text{M}$ |
| 33 | $k_{\text{ROS-MAFA}}$ | Rate of exclusion of MAFA from the nucleus by ROS | $0.06 \pm 0.015 \mu\text{M}$ |
| 34 | $k_{\text{GSK3-MAFA-pdeg}}$ | Rate of MAFA protein degradation by GSK3 | $0.006 \pm 0.002 \mu\text{M}$ |
| 35 | $V_{\text{mINS}}$ | Maximum rate of insulin mRNA production | $1.4 \pm 0.39 \mu\text{M}$ |
| 36 | $K_{\text{PDX1-INS}}$ | Dissociation constant of PDX1 from the binding site of insulin promoter | $2.34 \pm 0.47 \mu\text{M}$ |
| 37 | $K_{\text{MAFA-INS}}$ | Dissociation constant of MAFA from the binding site of insulin promoter | $2.29 \pm 0.47 \mu\text{M}$ |

### Validation

We validated our model against experimental data that were different from the data that we used for the parameter estimation. The details of the experimental data used for validation are given in the **Methods** section. The validation results are shown in Fig. 4. The validation was repeated for all 100 estimated parameter sets and the simulated values are presented as mean ± SD (n = 100). Fig. 4(a) and (b) show the comparison between experimental data and simulated values of the relative PDX1 and insulin mRNA expressions after 48 h and 72 h of exposure to high glucose concentration of 25 mM. Fig. 4(c) shows the relative PDX1 expression after 48 h exposure to 50 µM H_2_O_2_. Fig 4(d) shows the relative PDX1 expression in the presence of AKT inhibitor, GSK3 inhibitor, and both. Fig. 4(e) shows the relative MAFA levels after exposure to 50 µM H_2_O_2_ for 30, 60 and 90 minutes, and Fig. 4(f) shows the relative MAFA levels after exposure to 4 mM and 16.7 mM glucose.

**Figure 4:**
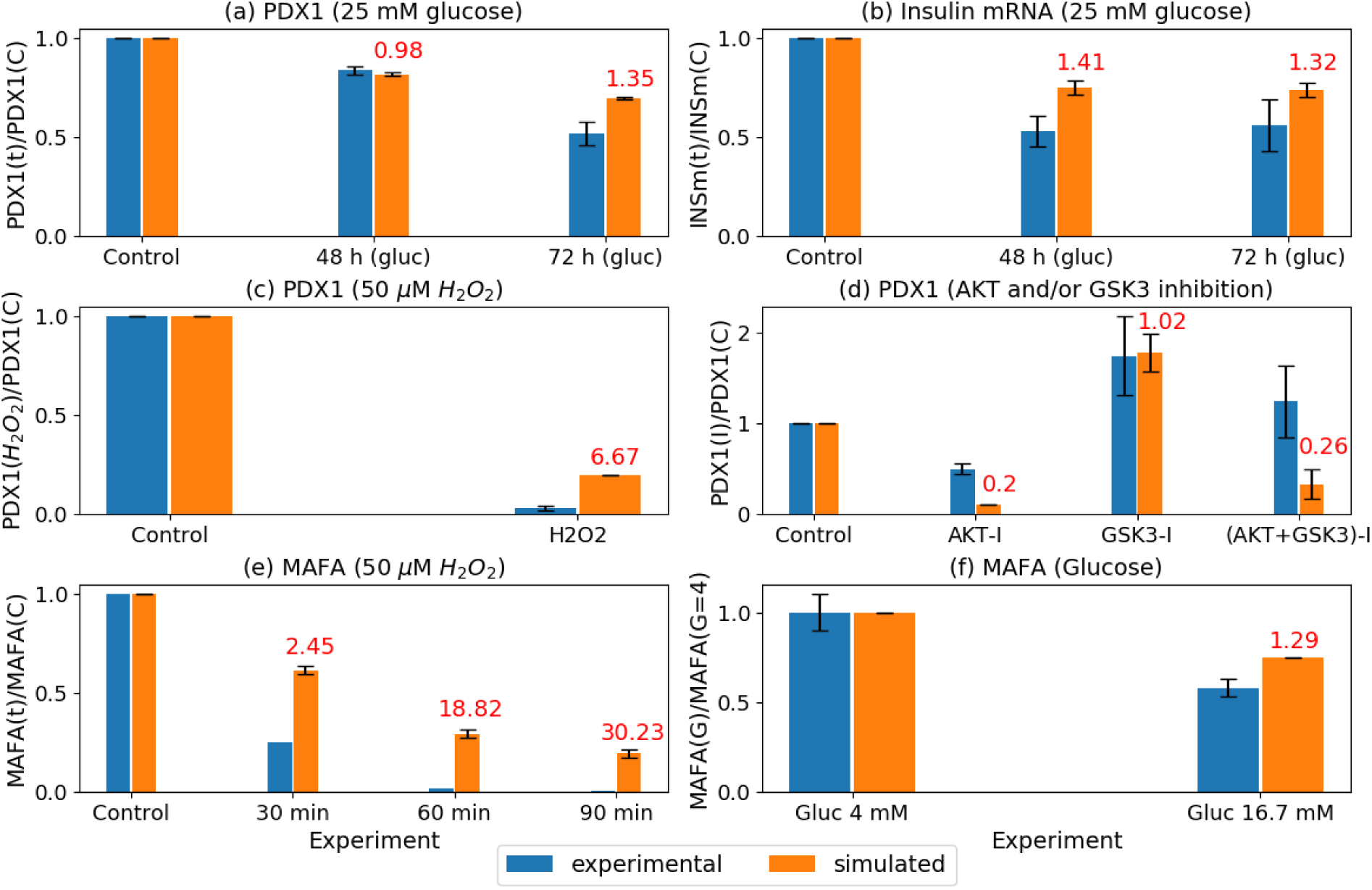
Validation: comparison between experimental data and simulated values of (a) relative PDX1 levels after 48 h and 72 h of exposure to high glucose concentration of 25 mM, (b) relative insulin mRNA levels after 48 h and 72 h of exposure to high glucose concentration of 25 mM, (c) relative PDX1 level after 48 h exposure to 50 µM H_2_O_2_, (d) relative PDX1 levels in the presence of AKT inhibitor, GSK3 inhibitor, and both, (e) relative MAFA levels after exposure to 50 µM H_2_O_2_ for 30, 60 and 90 minutes, and (f) relative MAFA levels after exposure to 4 mM and 16.7 mM glucose. All simulated values are presented as mean ± SD (n = 100). Here C denotes the Control case, I denotes the inhibited case and G denotes glucose. The values in red denote the fold-overestimation or underestimation compared to the corresponding experimental data.

Overall, our model simulations are successful in reproducing the trends observed from the experimental data. The model reproduces a decrease in PDX1 and insulin mRNA levels after 48 h and 72 h exposure to 25 mM glucose (Fig. 4(a) and (b)) and a decrease in PDX1 level after exposure to 50 µM H_2_O_2_ for 48 h (Fig. 4(c)). In Fig. 4(a), our model underestimates the PDX1 level by 0.98-fold and overestimates the PDX1 level by 1.35-fold compared to the experimental levels when exposed to 25 mM glucose for 48 h and 72 h respectively. In Fig. 4(b), we observe overestimations in insulin mRNA levels compared to the experimental data by 1.41-fold and 1.32-fold when exposed to 25 mM glucose for 48 h and 72 h respectively. In Fig. 4(c), our model overestimates the PDX1 level by 6.67-fold compared to the experimental data on exposure to 50 µM H_2_O_2_ for 48 h. In Fig. 4(d), compared to the experimental data, our model (i) underestimates PDX1 level by 0.2-fold in the presence of AKT inhibitor, (ii) overestimates PDX1 level by 1.02-fold in the presence of GSK3 inhibitor, and (iii) underestimates PDX1 level by 0.26-fold in the presence of both AKT and GSK3 inhibitors. The model also reproduces the decrease in MAFA levels after exposure to (i) 50 µM H_2_O_2_ for 30, 60 and 90 minutes (Fig. 4(e)) and (ii) 16.7 mM glucose (Fig. 4(f)). In Fig. 4(e), compared to the experimental data, our model overestimates the MAFA levels by 2.45-fold, 18.82-fold, and 30.23-fold when exposed to 50 µM H_2_O_2_ for 30, 60 and 90 minutes respectively. In Fig. 4(f), we observe a 1.29-fold overestimation in MAFA levels compared to the experimental data under exposure to 16.7 mM glucose for 48 h.

### Scenarios

The calibrated and validated mathematical model was then used to simulate the changes in steady-state levels of insulin mRNA, PDX1, MAFA, FOXO1, GSK3, JNKp, p38MAPKp and ROS (H_2_O_2_) at glucose concentrations of 3 mM, 10 mM, 20 mM and 30 mM (Fig. 5). All values reported in Fig. 5 are steady-state values. From Fig. 5 we observe that with increasing glucose concentrations, the steady-state levels of the transcription factors PDX1 and MAFA as well as insulin mRNA decrease. At the very high glucose concentration of 30 mM, their levels almost become negligible. On the other hand, the steady-state levels of FOXO1, GSK3, JNKp, p38MAPKp and ROS (H_2_O_2_) increase with increasing glucose concentrations. The high nuclear FOXO1 and GSK3 are associated with reduced expression of PDX1 and MAFA and consequently insulin mRNA.

**Figure 5:**
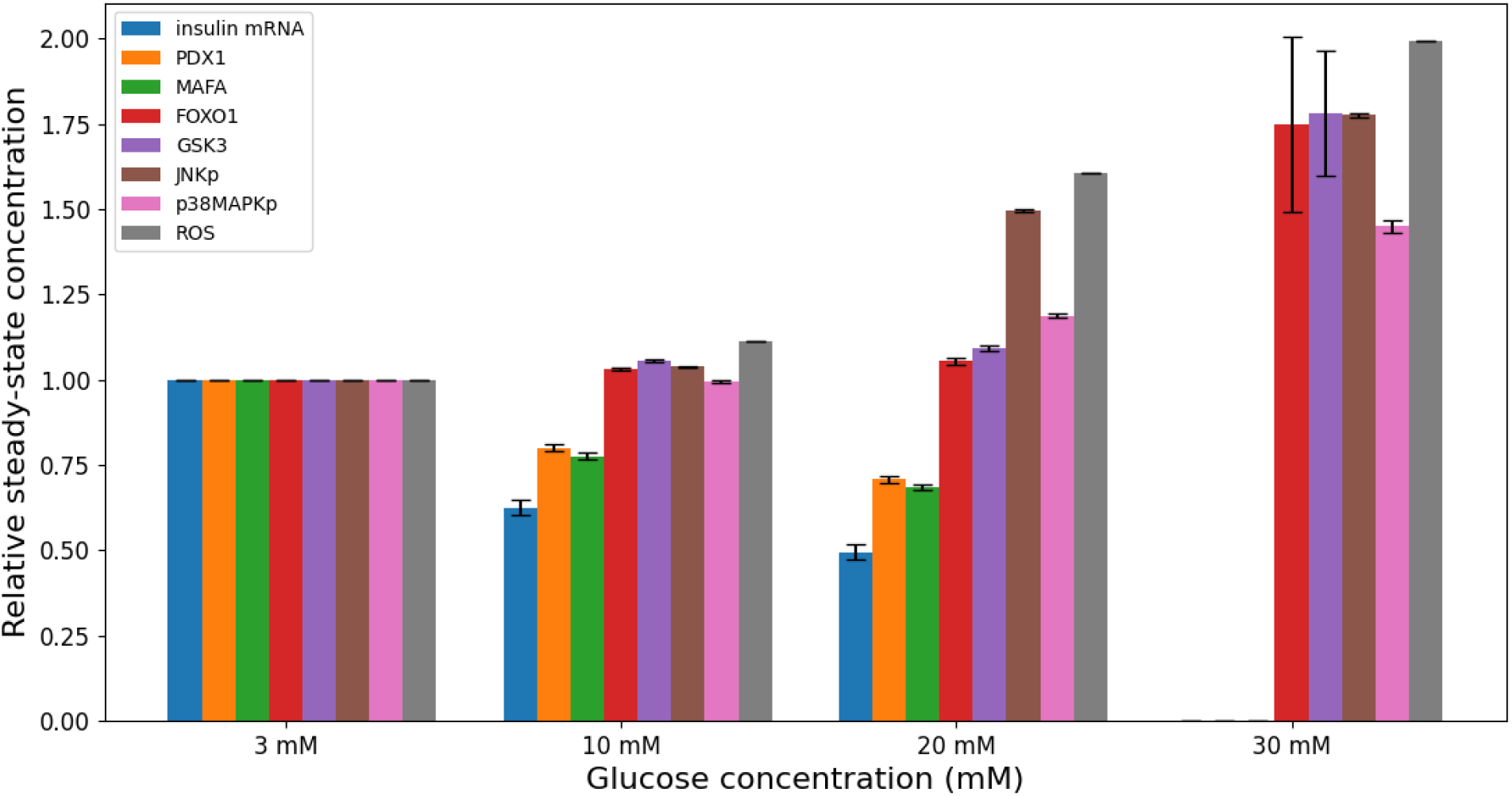
Simulated steady-state concentrations of insulin mRNA, PDX1, MAFA, nuclear FOXO1, nuclear GSK3, JNKp, p38MAPKp and ROS (H_2_O_2_) at glucose concentrations of 3 mM, 10 mM, 20 mM and 30 mM normalized by their steady-state concentrations at 3 mM glucose. All values are presented as mean ± SD (n = 100).

Furthermore, we used this model to simulate specific scenarios that could allow to restore PDX1 and MAFA levels and therefore insulin gene expression after exposure to high glucose levels (Fig. 6). We first simulated the steady-state levels of PDX1, MAFA and insulin mRNA by setting the glucose concentration to a high level of 25 mM. Next, we simulated the following scenarios to check the percentage restoration of PDX1, MAFA and insulin mRNA:

1. glucose concentration was decreased to normal glucose level of 5 mM,
2. glucose concentration was decreased to 5 mM and AKTp concentration was increased by 10% simultaneously,
3. glucose concentration was decreased to 5 mM and nuclear FOXO1 level was decreased by 10% simultaneously,
4. glucose concentration was decreased to 5 mM and nuclear GSK3 level was decreased by 10% simultaneously,
5. glucose concentration was decreased to 5 mM and nuclear FOXO1 and nuclear GSK3 levels were decreased by 10% simultaneously,
6. glucose concentration was decreased to 5 mM and JNKp level was decreased by 10% simultaneously,
7. glucose concentration was decreased to 5 mM and p38MAPKp level was decreased by 10% simultaneously, and
8. glucose concentration was decreased to 5 mM and nuclear FOXO1, JNKp and p38MAPKp levels were decreased by 10% simultaneously.

**Figure 6:**
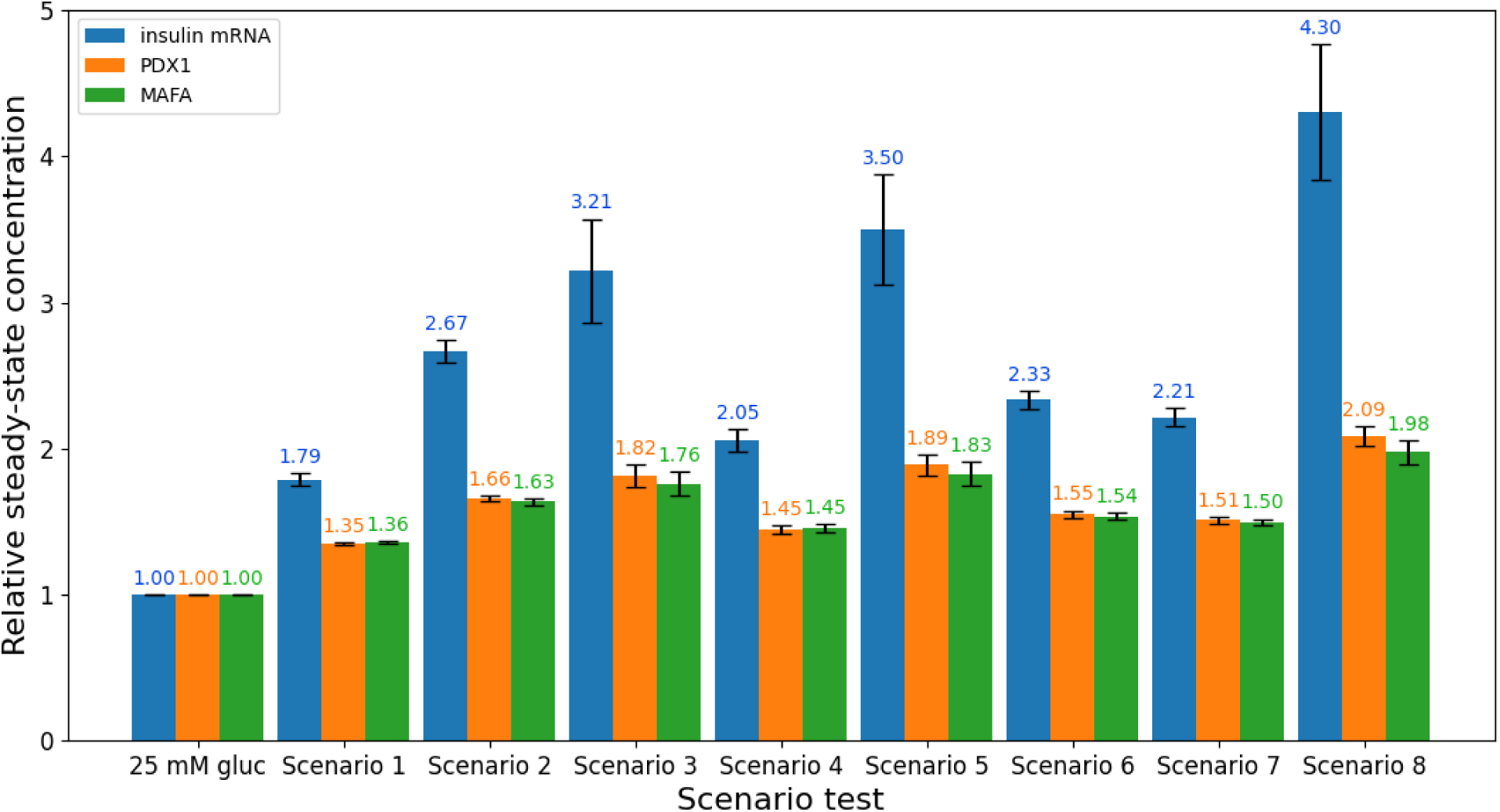
Eight scenarios that could allow to restore PDX1, MAFA and insulin mRNA levels after exposure to high glucose concentration of 25 mM. Scenario 1: glucose concentration was decreased to normal glucose level of 5 mM; Scenario 2: glucose concentration was decreased to 5 mM and AKTp concentration was increased by 10% simultaneously; Scenario 3: glucose concentration was decreased to 5 mM and nuclear FOXO1 level was decreased by 10% simultaneously; Scenario 4: glucose concentration was decreased to 5 mM and nuclear GSK3 level was decreased by 10% simultaneously; Scenario 5: glucose concentration was decreased to 5 mM and nuclear FOXO1 and nuclear GSK3 levels were decreased by 10% simultaneously; Scenario 6: glucose concentration was decreased to 5 mM and JNKp level was decreased by 10% simultaneously; Scenario 7: glucose concentration was decreased to 5 mM and p38MAPKp level was decreased by 10% simultaneously; Scenario 8: glucose concentration was decreased to 5 mM and nuclear FOXO1, JNKp, and p38MAPKp levels were decreased by 10% simultaneously. All values are steady-state, normalized by the steady-state levels of the respective variables at 25 mM glucose and are presented as mean ± SD (n = 100).

The resulting steady-state levels of PDX1, MAFA and insulin mRNA are given in Fig. 6. All reported steady-state concentrations in Fig. 6 were normalized by the steady-state concentration of the respective variables at 25 mM glucose. From the results, considering only changes in a single variable, we observe that a reduction in nuclear FOXO1 level (scenario 3) caused the highest increase in PDX1, MAFA and insulin mRNA levels (1.82, 1.76 and 3.21-fold respectively). Increasing AKTp concentration (scenario 2) resulted in the second highest increase in PDX1, MAFA and insulin mRNA levels, followed by the decrease in stress-activated kinases, JNKp and p38MAPKp (scenarios 6 and 7). Decrease in GSK3 level (scenario 4) was the least effective in restoring PDX1, MAFA and insulin mRNA levels.

Thus, from these scenario simulations, we see that regulating the concentration of nuclear FOXO1 has the highest effect on the steady-state levels of PDX1 and MAFA and thus on insulin mRNA. It has been shown in many studies that FOXO1 is an important transcription factor in the regulation of beta-cell identity transcription factors. FOXO1 has been in focus in recent studies since it plays a complex role in the pathophysiology of beta-cells and acts as a double-edged sword [52–54]. Since AKTp phosphorylates FOXO1 and prevents its translocation to the nucleus, increasing the concentration of AKTp also indirectly increases the steady-state levels of PDX1, MAFA and insulin mRNA. The stress-activated kinases, JNKp and p38MAPKp, facilitate the translocation of FOXO1 to the nucleus and JNKp inhibits the phosphorylation of AKT. Thus, decreasing the levels of these two kinases could also help to ameliorate the levels of PDX1, MAFA, and insulin mRNA after exposure to chronic hyperglycemia.

## Discussion

We developed an integrated mathematical model of the underlying mechanisms regulating two important beta-cell identity transcription factors and regulators of insulin promoter activity, PDX1 and MAFA, in the presence of chronic hyperglycemia. This model was able to reproduce experimentally measured levels of these transcription factors as well as insulin mRNA under different glucose concentrations. In addition, we simulated scenarios with different potential targets that could allow to restore PDX1 and MAFA levels after exposure to chronic hyperglycemia, where FOXO1 emerged as an important intervention target.

To develop this model, we integrated existing models of glucose transport, glucose metabolism, calcium dynamics, and protein folding from the literature and proposed new equations to describe the regulation of the beta-cell identity transcription factors, PDX1 and MAFA, by the insulin signaling pathway, stress-activated kinases (JNK and p38MAPK) and ROS (H_2_O_2_). We also developed new rate equations to describe the transcription of insulin gene by PDX1 and MAFA. To our knowledge there are no existing models of these processes in the literature. Thus, we built a model that integrates several models describing key processes underlying beta-cell function and links them with the pathways that control the beta-cell identity transcription factors. By doing so we propose a more comprehensive model of beta-cell (dys)-function and we connect these previously modelled processes to the mechanism of beta-cell loss of identity which is particularly important for the pathogenesis of T2D [54, 55].

We used parameter values from existing models as well as estimated some parameter values from experimental data. We performed the parameter estimation in two stages. The experimental data used for the first stage of parameter estimation have oscillations, whereas the simulated data do not (Fig. 2(a), (b), (c)). This is because the equations of the model do not include any oscillatory mechanisms themselves. Oscillations in glucose metabolism, membrane potential, and Ca^2+^ in the beta-cell due to increased glucose concentrations have been observed in many studies [56–58]. These oscillations are responsible for triggering insulin secretion [31, 59–61]. However, the aim of the current model is to study the pathways involved in the regulation of beta-cell identity transcription factors and insulin gene expression which are slower processes and hence not regulated by these oscillations. Thus, in the context of our model aim, we have not included the oscillatory mechanisms in our model equations. Another observed difference between the experimental data and simulated data is that with increasing glucose concentration the experimental data have a faster initial increase in NADH levels as compared to the initial increase obtained with the model (Fig. 2(a), (b), (c)). For instance, the experimental initial increase is approximately 1.8 and 2.5 times faster than the simulated initial increase at 10 mM and 20 mM glucose concentrations respectively ((Fig. 2(b), (c)). Since, we are only interested in the trends, we compare the steady-state values. In this regard, we observe that the levels reached by the simulated data are consistent with the experimental data. From Fig. 3(a), (b) and (e), we observe that the simulated data reach equilibrium, whereas the experimental data keep on decreasing. In our model, JNK is phosphorylated by H_2_O_2_ and UPR and is dephosphorylated at a constant rate. In reality, JNK and p38MAPK have other positive and negative regulators and these other regulators could be responsible for the behaviour of JNK and p38MAPK observed from the experimental data. For similar reasons, our model simulations could not reproduce the exact behaviour of phosphorylated AKT (AKTp) from the experimental data. Another possible reason could be that the experimental data are based on different cell lines and conditions. Using these experimental data together for the parameter estimation results in these discrepancies between the simulated behaviour and experimental behaviour. However, our model can reproduce the trends, that is, in the presence of high H_2_O_2_, the phosphorylation of JNK and p38MAPK increases, and in the presence of insulin stimulation the phosphorylation of AKT increases.

We validated the simulated levels of PDX1 and MAFA against a different set of experimental data. From the validation results, we observe that our model could closely reproduce the trends observed from experimental data. The discrepancies observed could be due to the fact that there are other pathways and factors which we have not included in our model. For example, in Fig. 4(d) we observe that when AKT is inhibited, the simulated level of PDX1 is much lower compared to the experimentally measured level. This could be because in our model, AKT is the only inhibitor of FOXO1 and GSK3, which are inhibitors of PDX1. Hence, in the absence of other suppressors, AKT inhibition greatly increases the levels of nuclear FOXO1 and GSK3 which causes the large decrease in PDX1 level. However, in reality, FOXO1 and GSK3 have other inhibitors. So, when AKT is inhibited in the experimental set-up, these other inhibitors prevent a large increase in the levels of nuclear FOXO1 and GSK3. Due to this reason, the levels in experimental and simulated PDX1 on AKT inhibition are different. In Fig. 4(c) and 4(e), the exposure to 50 µM H_2_O_2_ has a less pronounced effect on the downregulation of PDX1 and MAFA. From the stage 2 parameter estimation results (Fig. 3(a) and 3(b)), we observed that on average the level of phosphorylated JNK and p38MAPK increased less compared to the experimental data when exposed to 50 µM H_2_O_2_. Since these two proteins play an important role in downregulating PDX1 and MAFA, the less pronounced effect of H_2_O_2_ on the phosphorylation of JNK and p38MAPK could be a possible reason for the less pronounced effect of H_2_O_2_ on the downregulation of PDX1 and MAFA. However, despite these differences in the experimental and simulated levels, our model simulations could reproduce the experimentally observed trends.

We used this model to simulate specific scenarios where we regulated the levels of one or more proteins, namely AKTp, FOXO1, GSK3, JNKp, and p38MAPKp, to allow the restoration of PDX1, MAFA and insulin mRNA levels after exposure to high glucose concentration. From these simulations, we found that FOXO1 inhibition resulted in the highest restoration of PDX1, MAFA and insulin mRNA levels. In recent studies, FOXO1 has been identified as a potential transcription factor linking metabolic stress to beta-cell identity loss [54]. FOXO1 is a multifunctional transcription factor that plays both adaptive as well as deleterious roles in beta-cells under stress [54]. On the one hand, FOXO1 translocation to the nucleus has been associated with increased MAFA expression, whereas, on the other hand, FOXO1 translocation to the nucleus has been associated with inhibition of PDX1 transcription as well as nuclear exclusion of PDX1 protein. FOXO1, thus represents a potential therapeutic target in T2D and it is important to extensively model the interaction networks involving FOXO1.

It should be noted that this model is not exhaustive and may miss some relevant processes involved in the regulation of beta-cell transcription factors. For instance, cytokines and inflammation are important mediators of beta-cell dysfunction, which our model does not consider, and which can be a possible future extension to our model. Beta-cell dysfunction and compromised beta-cell identity result after years of chronic hyperglycemia. This long-term simulation is not possible with the proposed model. However, the model can be useful in simulating the trends that are observed in reality when glucose concentration is increased above physiological levels. In other words, the model simulations facilitate the study of the short-term mechanisms whose build-up leads to long-term beta-cell dysfunction and compromised beta-cell identity. Furthermore, this model can be scaled up from the single beta-cell level to the islet level to study the collective effect of compromised beta-cell identity in the presence of hyperglycemia.

Such computational models of regulation of key beta-cell identity transcription factors are important to develop specific therapies targeted towards reversal of compromised beta-cell identity. The important question is how beta-cell function can be restored after major loss of beta-cell identity transcription factors in T2D. Experiments are underway to identity pathways that could be targeted to reverse beta-cell identity loss [54, 62–64]. For example, [64] showed that a small molecule inhibitor of the transforming growth factor-beta receptor (ALK5) was able to reverse beta-cell identity loss in islets isolated from mice with extreme diabetes. They also showed that incubation of isolated human islets with ALK5 inhibitor resulted in increased expression of beta-cell identity markers, including insulin, PDX1 and MAFA. Thus, it is important to identify pathway-specific therapeutics to effectively control T2D and computational models like ours can help in this identification process. For example, we found through our model simulations that FOXO1 inhibition resulted in the highest restoration of PDX1, MAFA and insulin mRNA levels.

Our integrated mathematical model is a first step towards developing more holistic models which can further extend our understanding of the underlying non-linear biological mechanisms leading to compromised beta-cell identity and beta-cell dysfunction in the presence of chronic hyperglycemia. Our model can be extended by including other transcription factors associated with mature beta-cell identity such as NKX6.1 and NEUROD1. Such integrated computational models could help to identify the most important pathways and feedback loops given specific inputs and drivers, further understand the interaction between different pathways and processes, generate hypotheses and test possible intervention strategies.

## Methods

### Parameter estimation

Unknown parameters were estimated using the particle swarm optimization (PSO) algorithm [65] which can search very large spaces of candidate solutions and was therefore particularly adapted to our model which contains 37 free parameters. We implemented the PSO algorithm using the PySwarms toolkit [66].

The cost functions for the optimization were defined as the root mean squared error between the experimental data and the simulated data. The parameter estimation was done by considering all these experimental data simultaneously and the individual cost functions were added after normalizing them to be between 0 and 1. The details of the experimental data and the corresponding cost functions for the first and second stages of parameter estimation are given in Table 7 and Table 8 respectively.

**Table 7:** Details of the experimental data and the corresponding cost functions used for stage 1 of the parameter estimation.

| Experiment details | Reference | Cost function |
| --- | --- | --- |
| Steady-state values of NADH, mitochondrial membrane potential, cytosolic, mitochondrial, ER and extra-cellular calcium at sub-stimulatory glucose level of 3 mM. | [44-49, 51] | $C_{SS} = \sqrt{\frac{\sum_{t=0}^n \left( \frac{X_t^{\text{sim}}}{X^{\text{lit}}} - 1 \right)^2}{n+1}}$<br>$X_t^{\text{sim}}$: simulated value of state variable $X$ at time step $t$,<br>$X^{\text{lit}}$: steady-state value of state variable $X$ obtained from literature. |
| Ratio of NADH concentration at stimulatory glucose levels (8 mM, 10 mM, 20 mM) to NADH concentration at sub-stimulatory glucose level of 3mM.<br>Mouse pancreatic islets were used. Islet NADH responses to step glucose elevations from a baseline | [57] | $C_{\text{NADH}} = \sqrt{\frac{\sum_{t=0}^n \left( \frac{[NADH]_t^{\text{sim\_G8}}}{[NADH]_{\text{G3}}} - \text{ratio}_t^{\text{data\_G8}} \right)^2}{n+1}} + \sqrt{\frac{\sum_{t=0}^n \left( \frac{[NADH]_t^{\text{sim\_G10}}}{[NADH]_{\text{G3}}} - \text{ratio}_t^{\text{data\_G10}} \right)^2}{n+1}} + \sqrt{\frac{\sum_{t=0}^n \left( \frac{[NADH]_t^{\text{sim\_G20}}}{[NADH]_{\text{G3}}} - \text{ratio}_t^{\text{data\_G20}} \right)^2}{n+1}}$<br>$[NADH]_t^{\text{sim\_G8}}, [NADH]_t^{\text{sim\_G10}},$ |
| concentration of 3 mM to 8 mM, 10 mM and 20 mM were recorded. 15, 28 and 17 islet recordings were used respectively for the 3 cases. The data was recorded for a duration of 10 mins and the relative autofluorescence was recorded. |  | $[NADH]_t^{sim\_G20}$: simulated concentrations of NADH at time step $t$ in the presence of 8 mM, 10 mM and 20 mM glucose,<br>$[NADH]^{G3}$: steady-state concentration of NADH at 3 mM glucose,<br>$ratio_t^{data\_G8}, ratio_t^{data\_G10}, ratio_t^{data\_G20}$: ratio of NADH concentration at stimulatory glucose levels (8 mM, 10 mM, 20 mM) to NADH concentration at sub-stimulatory glucose level of 3 mM obtained from the experimental data. |
| Ratio of cytosolic ATP concentrations at stimulatory glucose levels of 8.3 mM, 16.7 mM and 25 mM to cytosolic ATP concentration at sub-stimulatory glucose level of 2.8 mM.<br>Mouse pancreatic islets were used. Relative fluorescence was recorded by increasing glucose concentration from a baseline concentration of 2.8 mM to 8.3 mM, 16.7 mM and | [67] | $C_{ATPC}$<br>$= \sqrt{\frac{\sum_{t=0}^n \left( \frac{[ATP_C]_t^{sim\_G8.3}}{[ATP_C]^{G3}} - ratio_t^{data\_G8.3} \right)^2}{n+1}}$ $+ \sqrt{\frac{\sum_{t=0}^n \left( \frac{[ATP_C]_t^{sim\_G16.7}}{[ATP_C]^{G3}} - ratio_t^{data\_G16.7} \right)^2}{n+1}}$ $+ \sqrt{\frac{\sum_{t=0}^n \left( \frac{[ATP_C]_t^{sim\_G25}}{[ATP_C]^{G3}} - ratio_t^{data\_G25} \right)^2}{n+1}}$<br>$[ATP_C]_t^{sim\_G8.3}, [ATP_C]_t^{sim\_G16.7}, [ATP_C]_t^{sim\_G25}$: simulated concentrations of cytosolic ATP at time step $t$ in the presence of 8.3 mM, 16.7 mM and 25 mM glucose, |
| 25 mM. The duration of the recording was 100 seconds and 8 islets were used. | | $[ATP_c]^{G3}$: steady-state concentration of cytosolic ATP at 3 mM glucose,<br>$ratio_t^{data\_G8.3}, ratio_t^{data\_G16.7}, ratio_t^{data\_G25}$: ratio of ATP concentration at stimulatory glucose levels (8 mM, 10 mM, 20 mM) to ATP concentration at sub-stimulatory glucose level of 3 mM obtained from the experimental data. |

**Table 8:**
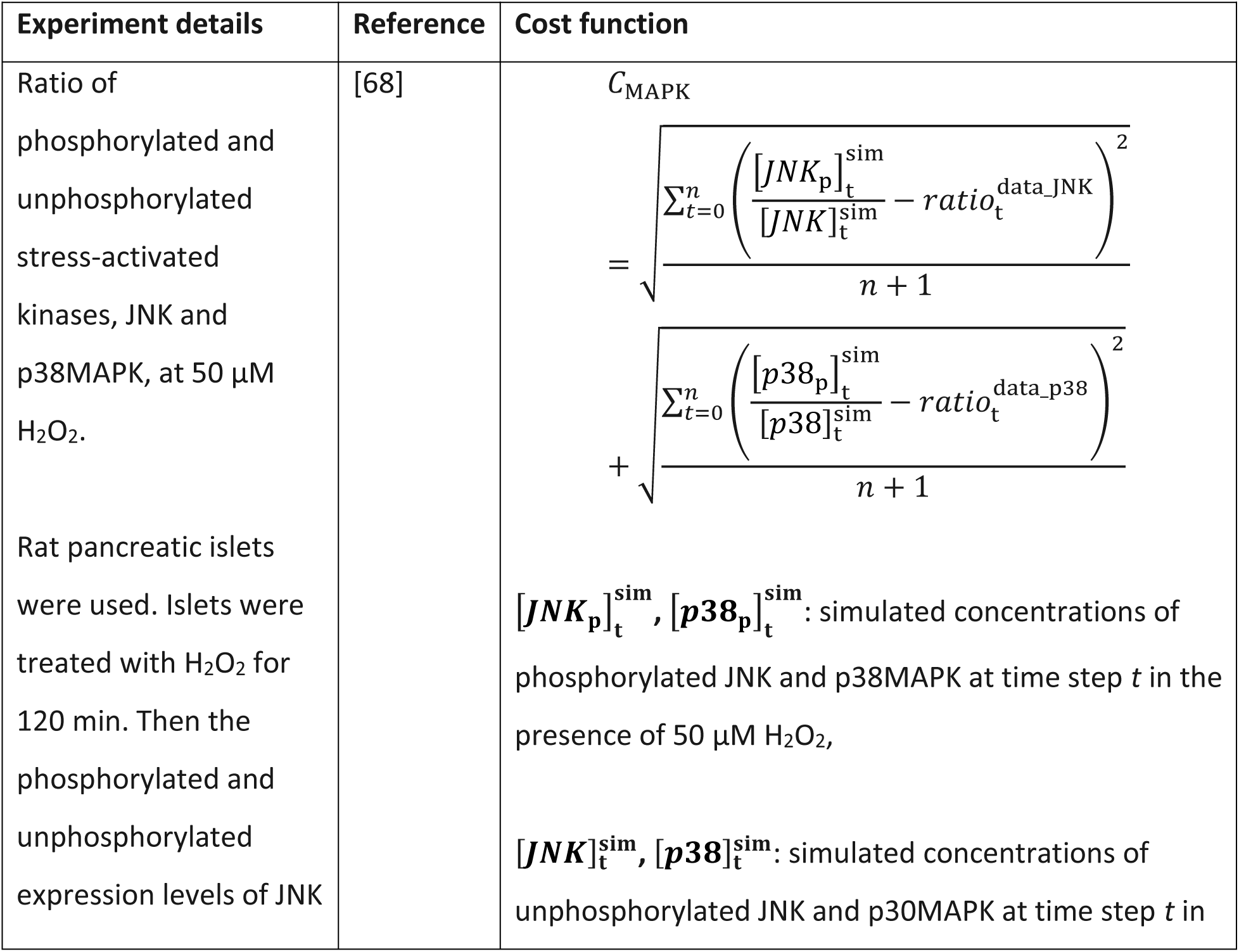
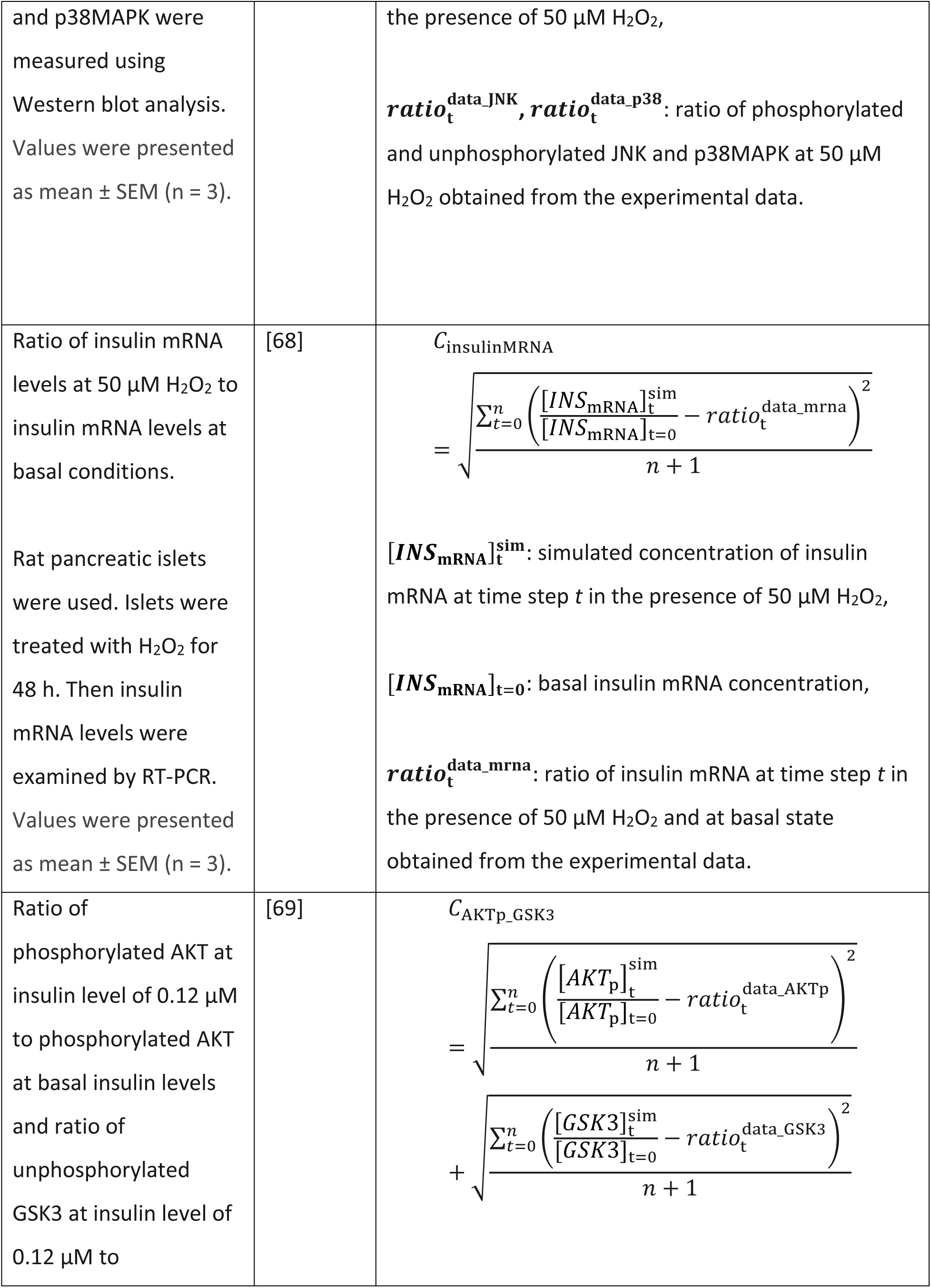
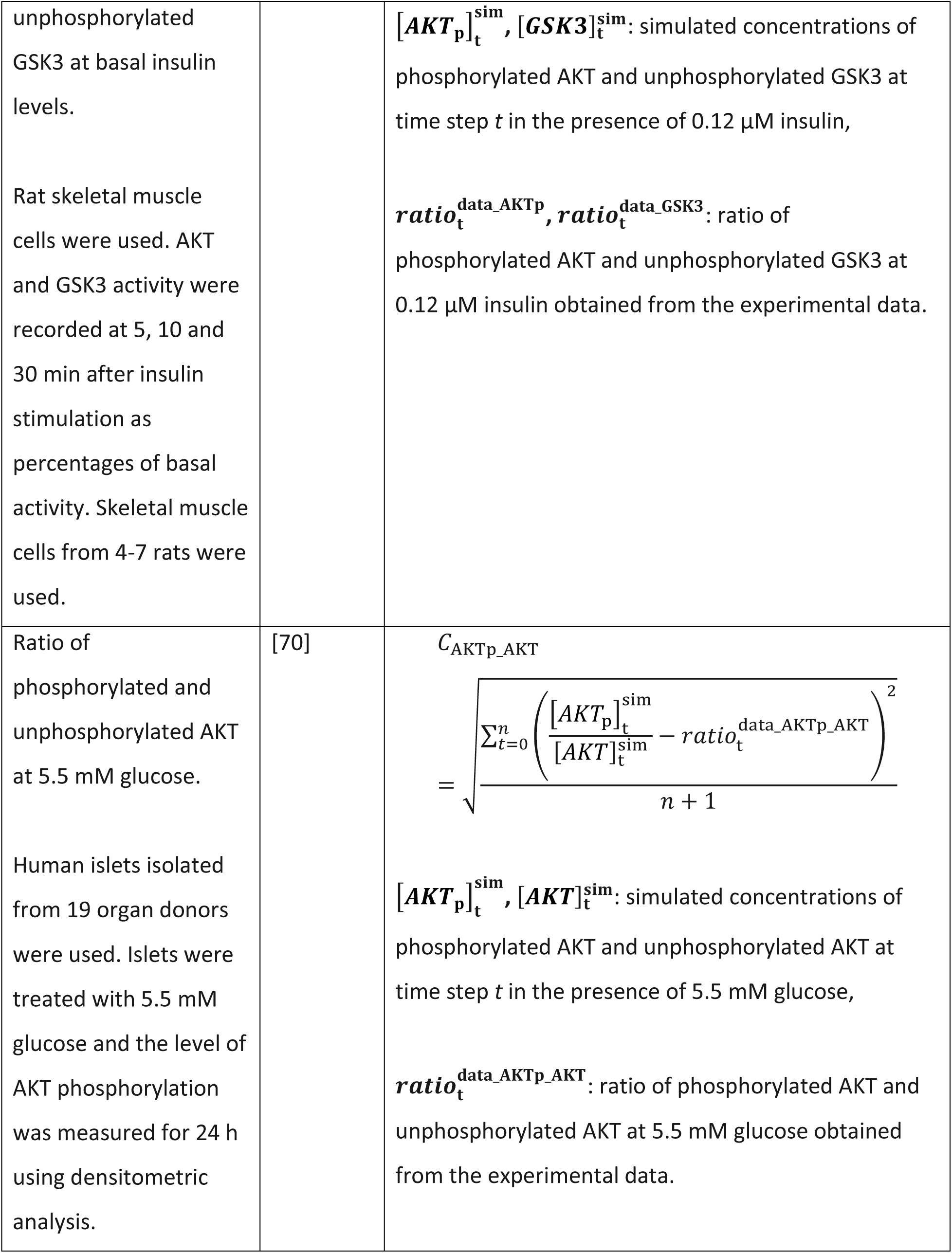
Details of the experimental data and the corresponding cost functions used for stage 2 of the parameter estimation.

| Experiment details | Reference | Cost function |
| --- | --- | --- |
| Ratio of phosphorylated and unphosphorylated stress-activated kinases, JNK and p38MAPK, at 50 μM H<sub>2</sub>O<sub>2</sub>.<br>Rat pancreatic islets were used. Islets were treated with H<sub>2</sub>O<sub>2</sub> for 120 min. Then the phosphorylated and unphosphorylated expression levels of JNK | [68] | $C_{MAPK}$ $= \sqrt{\frac{\sum_{t=0}^n \left( \frac{[JNK_p]_t^{sim}}{[JNK]_t^{sim}} - ratio_t^{data\_JNK} \right)^2}{n+1}} + \sqrt{\frac{\sum_{t=0}^n \left( \frac{[p38_p]_t^{sim}}{[p38]_t^{sim}} - ratio_t^{data\_p38} \right)^2}{n+1}}$<br>$[JNK_p]_t^{sim}, [p38_p]_t^{sim}$: simulated concentrations of phosphorylated JNK and p38MAPK at time step $t$ in the presence of 50 μM H<sub>2</sub>O<sub>2</sub>,<br>$[JNK]_t^{sim}, [p38]_t^{sim}$: simulated concentrations of unphosphorylated JNK and p30MAPK at time step $t$ in |
| and p38MAPK were measured using Western blot analysis. Values were presented as mean ± SEM (n = 3). | | the presence of 50 μM H<sub>2</sub>O<sub>2</sub>,<br>$ratio_t^{data\_JNK}, ratio_t^{data\_p38}$: ratio of phosphorylated and unphosphorylated JNK and p38MAPK at 50 μM H<sub>2</sub>O<sub>2</sub> obtained from the experimental data. |
| Ratio of insulin mRNA levels at 50 μM H<sub>2</sub>O<sub>2</sub> to insulin mRNA levels at basal conditions.<br>Rat pancreatic islets were used. Islets were treated with H<sub>2</sub>O<sub>2</sub> for 48 h. Then insulin mRNA levels were examined by RT-PCR. Values were presented as mean ± SEM (n = 3). | [68] | $C_{insulinMRNA}$ $= \sqrt{\frac{\sum_{t=0}^n \left( \frac{[INS_{mRNA}]_t^{sim}}{[INS_{mRNA}]_{t=0}} - ratio_t^{data\_mrna} \right)^2}{n + 1}}$<br>$[INS_{mRNA}]_t^{sim}$: simulated concentration of insulin mRNA at time step $t$ in the presence of 50 μM H<sub>2</sub>O<sub>2</sub>,<br>$[INS_{mRNA}]_{t=0}$: basal insulin mRNA concentration,<br>$ratio_t^{data\_mrna}$: ratio of insulin mRNA at time step $t$ in the presence of 50 μM H<sub>2</sub>O<sub>2</sub> and at basal state obtained from the experimental data. |
| Ratio of phosphorylated AKT at insulin level of 0.12 μM to phosphorylated AKT at basal insulin levels and ratio of unphosphorylated GSK3 at insulin level of 0.12 μM to | [69] | $C_{AKTp\_GSK3}$ $= \sqrt{\frac{\sum_{t=0}^n \left( \frac{[AKTp]_t^{sim}}{[AKTp]_{t=0}} - ratio_t^{data\_AKTp} \right)^2}{n + 1}}$ $+ \sqrt{\frac{\sum_{t=0}^n \left( \frac{[GSK3]_t^{sim}}{[GSK3]_{t=0}} - ratio_t^{data\_GSK3} \right)^2}{n + 1}}$ |
| unphosphorylated GSK3 at basal insulin levels.<br>Rat skeletal muscle cells were used. AKT and GSK3 activity were recorded at 5, 10 and 30 min after insulin stimulation as percentages of basal activity. Skeletal muscle cells from 4-7 rats were used. |  | $[AKT_p]_t^{sim}$, $[GSK3]_t^{sim}$: simulated concentrations of phosphorylated AKT and unphosphorylated GSK3 at time step $t$ in the presence of 0.12 μM insulin,<br>$ratio_t^{data\_AKTp}$, $ratio_t^{data\_GSK3}$: ratio of phosphorylated AKT and unphosphorylated GSK3 at 0.12 μM insulin obtained from the experimental data. |
| Ratio of phosphorylated and unphosphorylated AKT at 5.5 mM glucose.<br>Human islets isolated from 19 organ donors were used. Islets were treated with 5.5 mM glucose and the level of AKT phosphorylation was measured for 24 h using densitometric analysis. | [70] | $C_{AKTp\_AKT}$<br>$= \sqrt{\frac{\sum_{t=0}^n \left( \frac{[AKT_p]_t^{sim}}{[AKT]_t^{sim}} - ratio_t^{data\_AKTp\_AKT} \right)^2}{n + 1}}$<br>$[AKT_p]_t^{sim}$, $[AKT]_t^{sim}$: simulated concentrations of phosphorylated AKT and unphosphorylated AKT at time step $t$ in the presence of 5.5 mM glucose,<br>$ratio_t^{data\_AKTp\_AKT}$: ratio of phosphorylated AKT and unphosphorylated AKT at 5.5 mM glucose obtained from the experimental data. |

### Validation

The model was validated on experimental data that were different from those used for parameter estimation. The details of the experimental data used for validation ae given in Table 9.

**Table 9:** Experimental data used for validation.

| Details of experimental data | Reference |
| --- | --- |
| Relative PDX1 protein expression in 1.1B4 cells after 48 h and 72 h exposure to high glucose (25 mM). Values are mean ± SEM (n = 3). | [71] |
| Relative insulin mRNA expression in 1.1B4 cells after 48 h and 72 h exposure to high glucose (25 mM). Values are mean ± SEM (n = 3). | [71] |
| Oxidative stress–induced nucleo-cytoplasmic translocation of PDX1. HIT-T15 cells were used. Cells were exposed for 48 h to vehicle alone (control) and 50 μM H<sub>2</sub>O<sub>2</sub>. Values are mean ± SD (n = 3). | [72] |
| Densitometric analysis of PDX1 expression in human islets in the presence of vehicle (control), AKT inhibitor, GSK3 inhibitor, and both. Values are mean ± SD (n = 3). | [73] |
| Immunoblotting to determine MAFA levels in the nucleus of βTC-3 cells under exposure to 50 μM H<sub>2</sub>O<sub>2</sub> for 30, 60 and 90 minutes. | [6] |
| Relative MAFA levels in INS-1 cells treated for 48 h with 4 mM and 16.7 mM glucose (n=4). | [74] |

## Acknowledgements

We would like to thank Casper van Elteren (University of Amsterdam) for his help in developing the Cython code. This work is supported by the NTU Research Scholarship, ZonMw (Netherlands Organization for Health Research and Development, project number: 531003015), Social HealthGames (NWO, the Dutch Science Foundation, project number: 645.003.002), Computational Modelling of Criminal Networks and Value Chains (Nationale Politie, project number: 2454972), and TO-AITION (EU Horizon 2020 programme, call: H2020-SC1-2018-2020, grant number: 848146).

## Author contributions

PD, NM and RQ designed the model with biological insights from FC. PD developed the computational code and performed the simulations with the help of RQ. PD analyzed the results with the help of NM and RQ. PD and NM drafted the manuscript with critical inputs from RQ, FC and PMAS. All authors approved the final version.

