## Supplemental Material for "Compromised beta-cell identity in type 2 diabetes"

### 1 Model development

In the following sections we describe in detail all the important processes involved in beta-cell function, namely, glucose transport, glucose metabolism, calcium dynamics, electrical activity, insulin signaling, reactive oxygen species generation, insulin transcription, translation and folding, unfolded protein response and insulin secretion. We also explain the rate equations used to describe these processes.

#### 1.1 Rate equations for glucose transport

Glucose is transported into beta-cell by facilitated diffusion through plasma membrane-resident glucose transporter, GLUT-1. This glucose uptake through GLUT1 is modelled using Michaelis–Menten equations from Luni et al. [1].

Influx of extra-cellular glucose ($[Gluc]_{\text{e.c.}}$) through GLUT1 [1]:

$$J_{\text{G1I}} = \frac{V_{\text{mG1}}[Gluc]_{\text{e.c.}}}{K_{\text{mG1}} + [Gluc]_{\text{e.c.}}}. \qquad (S1)$$

Efflux of intra-cellular glucose ($[Gluc]_{\text{i.c.}}$) through GLUT1 [1]:

$$J_{\text{G1E}} = \frac{V_{\text{mG1}}[Gluc]_{\text{i.c.}}}{K_{\text{mG1}} + [Gluc]_{\text{i.c.}}} \qquad (S2)$$

where, $V_{\text{mG1}}$ is the maximum reaction rate for GLUT1 that is observed at saturating glucose concentrations, and $K_{\text{mG1}}$ is the glucose concentration at which the reaction rate is half of $V_{\text{mG1}}$.

**1.2 Rate equations for glucose metabolism**

Glucose is metabolized to produce energy in the form of adenosine triphosphate (ATP). The production of ATP from glucose is regulated by 3 main processes: glycolysis (in the cytosol), tricarboxylic acid (TCA) cycle (in the mitochondria) and electron transport chain (ETC) (in the mitochondria) [2].

Glycolysis is modelled by the activity of glucokinase which is described by the rate equation [3]:

$$J_{\text{GK}} = V_{\text{mGK}} \left( \frac{[Gluc]_{\text{i.c.}}^{n_{\text{GK}}}}{K_{\text{GK}}^{n_{\text{GK}}} + [Gluc]_{\text{i.c.}}^{n_{\text{GK}}}} \right) \left( \frac{[ATP]_{\text{C}}}{K_{\text{mATP}} + [ATP]_{\text{C}}} \right) \qquad (S3)$$

where, $[ATP]_{\text{C}}$ denotes the concentration of cytosolic ATP, $V_{\text{mGK}}$ is the maximum rate of glucose consumption, $K_{\text{mATP}}$ is the Michaelis-Menten constant, $K_{\text{GK}}$ is the glucose concentration at which the reaction rate is half of $V_{\text{mGK}}$, and $n_{\text{GK}}$ is the Hill coefficient.

Pyruvate is the main end-product of glycolysis and it enters the mitochondria for supply to the TCA cycle [2]. The TCA cycle is modelled by the activity of pyruvate dehydrogenase (PDH) which is described by the rate equation [3, 4]:

$$J_{\mathrm{PDH}} = V_{\mathrm{mPDH}} \left( \frac{[Pyr]}{K_{\mathrm{mPyr}} + [Pyr]} \right) \left( \frac{1}{q_1 + [NADH]/[NAD]} \right) \left( \frac{[Ca^{2+}]_{\mathrm{M}}}{q_2 + [Ca^{2+}]_{\mathrm{M}}} \right) \qquad (S4)$$

where, $[Pyr]$ denotes the concentration of pyruvate, $[Ca^{2+}]_{\mathrm{M}}$ denotes the concentration of mitochondrial calcium, $V_{\mathrm{mPDH}}$ is the maximum rate of PDH reaction, $K_{\mathrm{mPyr}}$ is the Michaelis-Menten constant, $q_1$ is the Michaelis-Menten-like constant for reduction of $NAD^+$ to NADH by the TCA cycle, and $q_2$ is the $S_{0.5}$ value for activation the TCA cycle by $[Ca^{2+}]_{\mathrm{M}}$. The calcium-dependent factor reflects the activation of PDH by $Ca^{2+}$.

The TCA cycle generates the reducing equivalents, NADH and $FADH_2$, which are transferred to the ETC for ATP synthesis [2]. The ETC is a collection of membrane-embedded proteins and organic molecules organized into four large complexes labeled I to IV. Complex I oxidizes NADH to generate two electrons. As electrons move through complex I in a series of redox reactions, energy is released, and the complex uses this energy to pump protons from the matrix into the intermembrane space of the mitochondria. $FADH_2$ is oxidized by complex II. Complex II does not pump protons across the membrane. In this model, we only consider complex I, since NADH is good at electron donation whereas $FADH_2$ is not. Also, complex I pumps protons across the membrane whereas complex II does not. The transport chain builds a proton gradient across the inner mitochondrial membrane, with a higher concentration of $H^+$ in the intermembrane space and a lower concentration in the matrix. As $H^+$ ions flow down their gradient and back into the matrix, they pass through an enzyme called ATP synthase ($F_1F_0$ATPase), which harnesses the flow of protons to synthesize ATP.

The flux, $J_{\mathrm{O}}$ [4] represents the rate at which NADH is oxidized and the rate at which $H^+$ are pumped by complex I. The exponential factor takes into account the change of this flux with variation in the proton concentration gradient. That is, the pumping of protons leads to an increase in the mitochondrial membrane potential ($\Psi_{\mathrm{M}}$).

$$J_{\mathrm{O}} = V_{\mathrm{O}} \left( \frac{[NADH]}{q_3 + [NADH]} \right) \left( 1 + \mathrm{e}^{\Psi_{\mathrm{M}} - q_4 / q_5} \right)^{-1} \qquad (S5)$$

where, $[NADH]$ denotes the concentration of mitochondrial NADH, $V_{\mathrm{O}}$ is the maximum rate of NADH oxidation by ETC, $q_3$ is the Michaelis-Menten constant for NADH consumption by the ETC, $q_4$ is the voltage dependence coefficient 1 of ETC activity, and $q_5$ is the voltage dependence coefficient 2 of ETC activity.

The rate of ATP synthesis by the mitochondrial $F_1F_0$ATPase is described by the rate equation [4, 5]:

$$J_{\mathrm{FIFO}} = V_{\mathrm{FIFO}}\left(\frac{q_6}{q_6 + [ATP]_{\mathrm{M}}}\right)\left(1 + \mathrm{e}^{q_7 - \Psi_{\mathrm{M}}/q_8}\right)^{-1} \quad (S6)$$

where, $[ATP]_{\mathrm{M}}$ denotes the concentration of mitochondrial ATP, $V_{\mathrm{FIFO}}$ is the maximum rate $F_1F_0$ATPase, $q_6$ is the inhibition constant of ATPase activity by ATP, $q_7$ is the voltage dependence coefficient 1 of ATPase activity, and $q_8$ is the voltage dependence coefficient 2 of ATPase activity. ATP synthesis is driven by the proton flux from the cytosol into the mitochondria, which depolarizes the mitochondrial membrane. Thus, $J_{\mathrm{FIFO}}$ has a steep sigmoidal dependency on mitochondrial membrane potential.

The exchange flux of $[ADP]_{\mathrm{C}}$ and $[ATP]_{\mathrm{M}}$ across mitochondrial membrane through adenine nucleotide translocator (ANT) is defined as [4]:

$$J_{\mathrm{ANT}} = V_{\mathrm{ANT}}\left(\frac{1 - \frac{\alpha_{\mathrm{C}}}{\alpha_{\mathrm{M}}}\frac{[ATP]_{\mathrm{C}}}{[ADP]_{\mathrm{C}}}\frac{[ADP]_{\mathrm{M}}}{[ATP]_{\mathrm{M}}}\mathrm{e}^{-F'\Psi_{\mathrm{M}}}}{\left(1 + \alpha_{\mathrm{C}}\frac{[ATP]_{\mathrm{C}}}{[ADP]_{\mathrm{C}}}\mathrm{e}^{-0.5\times F'\Psi_{\mathrm{M}}}\right)\left(1 + \frac{1}{\alpha_{\mathrm{M}}}\frac{[ADP]_{\mathrm{M}}}{[ATP]_{\mathrm{M}}}\right)}\right) \quad (S7)$$

where, $[ADP]_{\mathrm{C}}$ and $[ADP]_{\mathrm{M}}$ denote the concentrations of cytosolic and mitochondrial ADP respectively, $V_{\mathrm{ANT}}$ is the maximum rate of the ANT, $\alpha_{\mathrm{C}}$ and $\alpha_{\mathrm{M}}$ are factors taking cytosolic ADP and ATP buffering, and mitochondrial ADP and ATP buffering respectively into account, and $F' = \frac{F}{R\times T}$, where F is the Faraday constant, *R* is the perfect gas constant and *T* is the temperature.

The rate equation for ATP hydrolysis:

$$J_{\mathrm{HYD}} = \frac{J_{\mathrm{SERCA}}}{2} + k_{\mathrm{HYD}} \times [ATP]_{\mathrm{C}} \quad (S8)$$

where, $k_{\mathrm{HYD}}$ is the maximal rate of ATP hydrolysis and $K_{\mathrm{h}}$ is the Michaelis-Menten constant for ATP hydrolysis. The first term of the equation taken from [4] denotes the hydrolysis of ATP required to provide energy to the ER SERCA pumps to transport $Ca^{2+}$ into the ER. $J_{\mathrm{SERCA}}$ is the flux of $Ca^{2+}$ from cytosol into ER through SERCA and is described in the section 1.4: **Calcium dynamics**. The second term represents all other ATP-consuming processes in the cytosol.

### 1.3 Rate equations for electrical activity

When stimulated with glucose, beta-cells exhibit repetitive, bursting electrical activity. These oscillations in the beta-cell are caused by voltage-gated $K^+$ channels, voltage-gated calcium channels, K-ATP channels modulated by cytoplasmic ATP/ADP ratio, and glucose-dependent cytoplasmic changes of intracellular calcium concentration.

The conductance, $g_{\mathrm{K}}$, and current, $\mathrm{I_K}$, through the voltage-gated $K^+$ channels is described as [6, 7]:

$$g_{\mathrm{K}} = \overline{g_{\mathrm{K}}} \times n^4 \qquad (S9)$$

$$I_{\mathrm{K}} = g_{\mathrm{K}} \times \left(V_{\mathrm{K}} - V_{\mathrm{p}}\right) \qquad (S10)$$

where, $\overline{g_{\mathrm{K}}}$ is the maximum conductance per cell and $n$ is the fraction of activated $K^+$ channel. The activation of $K^+$ channels, *n*, is given as [6, 7]:

$$\frac{dn}{dt} = \frac{n_{\mathrm{inf}} - n}{\tau_{\mathrm{n}}} \qquad (S11)$$

where, $n_{\mathrm{inf}}$ is the steady-state value of *n* and is determined by the following equation [6, 7]:

$$n_{\mathrm{inf}} = \frac{1}{1 + e^{(V_{\mathrm{n}} - V_{\mathrm{p}})/S_{\mathrm{n}}}}. \qquad (S12)$$

The conductance, $g_{\mathrm{Ca}}$, and current, $I_{\mathrm{Ca}}$, through the voltage-gated calcium channels is described as [6, 7]:

$$g_{\mathrm{Ca}} = \overline{g_{\mathrm{Ca}}} \times \left(P_{\mathrm{O}} \times X_{\mathrm{f}} + m_{\mathrm{infs}} \times J \times (1 - X_{\mathrm{f}})\right) \qquad (S13)$$

$$I_{\mathrm{Ca}} = g_{\mathrm{Ca}} \times g_{\mathrm{hk}} \qquad (S14)$$

where, $g_{\mathrm{hk}} = [Ca^{2+}]_{\mathrm{e.c.}} \times \frac{V_{\mathrm{p}}}{1-e^{V_{\mathrm{p}}/RT\_2F}}$.

In the above equations, $\overline{g_{\mathrm{Ca}}}$ is the maximum conductance per cell and $X_{\mathrm{f}}$ is a fraction of the maximum conductance. $\mathrm{P_O}$ denotes the fraction of open fast channels and is defined as [6, 7]:

$$\frac{dP_{\mathrm{O}}}{dt} = k_{\mathrm{neg}} \times (1 - P_{\mathrm{O}} - P_{\mathrm{C}}) - k_{\mathrm{star}} \times g_{\mathrm{hk}} \times P_{\mathrm{O}} + \alpha \times P_{\mathrm{C}} - \beta \times P_{\mathrm{O}} \qquad (S15)$$

where, $P_{\mathrm{C}}$ is the fraction of closed fast channels defined as [6, 7]:

$$\frac{dP_{\mathrm{C}}}{dt} = -\alpha \times P_{\mathrm{C}} + \beta \times P_{\mathrm{O}} \qquad (S16)$$

where, $\alpha = \frac{m_{\mathrm{inff}}}{\tau_{\mathrm{m}}}$ and $\beta = \frac{1-m_{\mathrm{inff}}}{\tau_{\mathrm{m}}}$. $m_{inff}$ denotes the voltage-dependent activation of the fast channels and is defined as [6, 7]:

$$m_{\mathrm{inff}} = \frac{1}{1 + e^{(V_{\mathrm{mf}}-V_{\mathrm{p}})/S_{\mathrm{mf}}}}. \qquad (S17)$$

*J* is the fraction of activated slow channels defined as [6, 7]:

$$\frac{dJ}{dt} = \frac{J_{\mathrm{inf}} - J}{\tau_{\mathrm{J}}} \qquad (S18)$$

where, $J_{\mathrm{inf}}$ is the steady-state value of *J* and is determined by the following equation [6, 7]:

$$J_{\mathrm{inf}} = \frac{1}{1 + e^{(V_{\mathrm{p}}-V_{\mathrm{J}})/S_{\mathrm{J}}}}. \qquad (S19)$$

$m_{\mathrm{infs}}$ denotes the voltage-dependent activation of the slow channels and is defined as [6, 7]:

$$m_{\mathrm{infs}} = \frac{1}{1 + e^{(V_{\mathrm{ms}}-V_{\mathrm{p}})/S_{\mathrm{ms}}}}. \qquad (S20)$$

ATP produced from glucose metabolism in the mitochondria is released into the cytosol which increases the ATP/ADP ratio and closes the K-ATP channel on the plasma membrane causing depolarization [2]. This facilitates the entry of $Ca^{2+}$ into the cytosol [2].

The conductance, $g_{\mathrm{K-ATP}}$, and current, $I_{\mathrm{K-ATP}}$, through the K-ATP channels modulated by cytosolic ATP/ADP ratio is described as [7]:

$$g_{\mathrm{K-ATP}} = \overline{g_{\mathrm{K-ATP}}} \times \frac{1 + \frac{[ADP]_{\mathrm{C}}}{K_{\mathrm{ADP-bind}}}}{1 + \frac{[ADP]_{\mathrm{C}}}{K_{\mathrm{ADP-bind}}} + \frac{[ATP]_{\mathrm{C}}}{K_{\mathrm{ATP-bind}}}} \qquad (S21)$$

$$I_{\mathrm{K-ATP}} = g_{\mathrm{K-ATP}} \times \left(V_{\mathrm{K}} - V_{\mathrm{p}}\right) \qquad (S22)$$

where, $\overline{g_{\mathrm{K-ATP}}}$ is the maximum conductance per cell, $K_{\mathrm{ADP-bind}}$ is the ADP binding constant, and $K_{\mathrm{ATP-bind}}$ is the ATP binding constant.

### 1.4 Calcium dynamics

The cytosolic $Ca^{2+}$ concentration regulates a vast number of processes, including metabolism, transcription, and secretion. The cytosolic $Ca^{2+}$ concentration is maintained by the bi-directional movement of $Ca^{2+}$ between cytosol and extra-cellular, mitochondria and ER.

#### 1.4.1 Calcium handling by mitochondria

The flux of $Ca^{2+}$ from cytosol into mitochondria mediated by the mitochondrial $Ca^{2+}$ uniporter (MCU) is defined as [4]:

$$J_{\mathrm{MCU}} = V_{\mathrm{MCU}} \times \left(\frac{[Ca^{2+}]_{\mathrm{C}}^2}{K_{\mathrm{MCU}}^2 + [Ca^{2+}]_{\mathrm{C}}^2}\right) e^{p_1 \times \Psi_{\mathrm{M}}} \qquad (S23)$$

where, $V_{\mathrm{MCU}}$ is the maximum $Ca^{2+}$ flux through MCU, $K_{\mathrm{MCU}}$ is the half-maximal activation constant for MCU, and $p_1$ is the voltage dependence coefficient of MCU activity. This flux is cooperatively stimulated by cytosolic $Ca^{2+}$ through the MCU1 subunit and increases exponentially with the potential difference across the mitochondrial membrane.

The flux of $Ca^{2+}$ from mitochondria to cytosol through the mitochondrial $Na^+/Ca^{2+}$ exchanger (NCX) is defined as [4]:

$$J_{\mathrm{NCX}} = V_{\mathrm{NCX}} \times \left(\frac{[Ca^{2+}]_{\mathrm{M}}}{[Ca^{2+}]_{\mathrm{C}}}\right) e^{p_2 \times \Psi_{\mathrm{M}}} \qquad (S24)$$

where, $V_{\mathrm{NCX}}$ is the maximum $Ca^{2+}$ flux through NCX and $p_2$ is the voltage dependence coefficient of NCX activity. This channel exchanges 1 $Ca^{2+}$ for 3 $Na^{+}$. It is assumed that the cytosolic $Na^{+}$ concentration remains constant and that channel activity is favoured by a large ratio of $Ca^{2+}$ concentrations between the cytosol and mitochondria.

#### 1.4.2 Calcium handling by ER

The flux of $Ca^{2+}$ from cytosol into ER through the unidirectional sarco/endoplasmic reticulum $Ca^{2+}$ ATPase (SERCA) is defined as [4],

$$J_{\mathrm{SERCA}} = V_{\mathrm{SERCA}} \times \left(\frac{[Ca^{2+}]_{\mathrm{C}}^2}{K_{\mathrm{p}}^2 + [Ca^{2+}]_{\mathrm{C}}^2}\right)\left(\frac{[ATP]_{\mathrm{C}}}{K_{\mathrm{e}} + [ATP]_{\mathrm{C}}}\right) \qquad (S25)$$

where, $V_{\mathrm{SERCA}}$ is the maximum $Ca^{2+}$ flux through SERCA, $K_{\mathrm{p}}$ is the dissociation constant of $Ca^{2+}$ from SERCA, and $K_{\mathrm{e}}$ is the dissociation constant of ATP from SERCA. The SERCA transports $Ca^{2+}$ from the cytosol into the ER, against the concentration gradient, using the energy provided by the hydrolysis of ATP.

The flux of $Ca^{2+}$ from ER to cytosol through inositol trisphosphate receptor ($IP_3R$) is defined as [4],

$$J_{\mathrm{IP3R}} = V_{\mathrm{IP3R}} \times IR_{\mathrm{a}} \times ([Ca^{2+}]_{\mathrm{ER}} - [Ca^{2+}]_{\mathrm{C}}) \qquad (S26)$$

where, $[Ca^{2+}]_{\mathrm{ER}}$ denotes the concentration of calcium in the ER, $V_{\mathrm{IP3R}}$ is the maximum $Ca^{2+}$ flux through $IP_3R$, and $IR_a$ is the fraction of active (open) $IP_3$ receptors given by,

$$IR_{\mathrm{a}} = 0.5 \times (1 - R_{\mathrm{i}}) \times \frac{IP3^3}{K_{\mathrm{IP3}}^3 + IP3^3} \qquad (S27)$$

where, $R_{\mathrm{i}}$ is the fraction of inactive receptors which changes with $Ca^{2+}$ as [4]:

$$\frac{dR_i}{dt} = k_{\mathrm{pos}} \times [Ca^{2+}]_{\mathrm{C}}^{ni} \times \frac{(1 - R_i)}{1 + \left({[Ca^{2+}]_{\mathrm{C}}}/{K_a}\right)^{na}} - k_{\mathrm{neg}} \times R_i. \qquad (S28)$$

#### 1.4.3 Calcium handling by cytosol

The $Ca^{2+}$ flux across the plasma membrane, where $Ca^{2+}$ enters the cytosol through the voltage-gated calcium channels and is removed from the cytosol through the plasma membrane $Ca^{2+}$ ATPase (PMCA) is defined as [7, 8]:

$$J_{\mathrm{PM}} = -\gamma \times I_{\mathrm{Ca}} - k_{\mathrm{PMCa}} \times [Ca^{2+}]_{\mathrm{C}} \qquad (S29)$$

where, $\gamma$ converts fA to $\mu M.s^{-1}$, and $k_{\mathrm{PMCa}}$ is the rate constant for the removal of $Ca^{2+}$ through PMCA.

**1.5 Reactive oxygen species (ROS)**

The mitochondrial electron transport chain [9, 10], and the ER protein-folding machinery [11] are two major sources of reactive oxygen species (ROS). Under physiological conditions, 0.2-2% of the electrons in the ETC do not follow the normal transfer order but instead directly leak out of the ETC and interact with oxygen to produce superoxide [12]. The superoxide undergoes dismutation by the superoxide dismutase, MnSOD, to form hydrogen peroxide ($H_2O_2$) [12].

Superoxide ($R_{\mathrm{s}}$) is generated by mitochondrial ETC and is converted to $H_2O_2$ ($R_{\mathrm{h}}$) by MnSOD ($A_{\mathrm{s}}$). The rate of production of superoxide is described by:

$$v_{\mathrm{superoxide-prod}} = q_{\mathrm{r}} \times J_{\mathrm{O}} \qquad (S30)$$

and the rate of conversion of superoxide to $H_2O_2$ by MnSOD is described by [13]:

$$v_{\mathrm{superoxide-to-H2O2}} = k_{\mathrm{rs}} \times R_{\mathrm{s}} \times A_{\mathrm{s}} \qquad (A31)$$

where, $q_{\mathrm{r}}$ is the fractional superoxide production and $k_{\mathrm{rs}}$ is the superoxide dismutation rate. The rate of clearance of $H_2O_2$ by catalase ($A_{\mathrm{h}}$) is defined as [13]:

$$v_{\mathrm{H2O2-clearance}} = \delta_{\mathrm{rh}} \times R_{\mathrm{h}} \times A_{\mathrm{h}} \qquad (S32)$$

where, $\delta_{\mathrm{rh}}$ is the rate $H_2O_2$ removal by catalase.

The protein folding process also contributes to the production of $H_2O_2$. The proinsulin molecule contains three disulphide bonds and the formation of each disulphide bond involves the disposal via oxidative pathways of two electrons giving rise to a molecule of hydrogen peroxide ($H_2O_2$) [11]. The rate of $H_2O_2$ generation by ER is described as:

$$v_{\mathrm{H2O2-prod-ER}} = 3 \times v_{\mathrm{folding-deg-chap}} \qquad (S33)$$

where, $v_{\mathrm{folding-deg-chap}}$ is the rate of protein folding and degradation which is described in detail in the section **1.7: Protein folding and unfolded protein response.**

### 1.6 Insulin signaling pathway, stress-activated kinases, and transcription factors

The transcription factors, PDX1 and MAFA, are important transcription factors for insulin gene transcription [14]. In normal conditions, MAFA is localized in the nucleus and PDX1 is shuttled between the cytosol and nucleus [14]. However, prolonged hyperglycemia impairs insulin gene expression by diminishing the expression and binding activity of PDX1 and MAFA to the insulin gene promoter. The signaling pathways mediating inhibition of insulin gene expression by oxidative stress and ER stress involve stress-activated kinases, c-Jun N-terminal kinase (JNK) and p38 mitogen-activated protein kinase (p38MAPK).

Glucose stimulates insulin secretion which results in the autocrine stimulation of the insulin receptors in the beta-cell and the insulin signaling pathway is activated. This activates protein kinase B (PKB aka AKT) which phosphorylates forkhead box protein O1 (FOXO1) and keeps it in the cytosol [15, 16]. If FOXO1 translocates to the nucleus it exerts its repressive action on PDX1 by two ways: downregulating PDX1 transcription and causing nuclear exclusion of PDX1 [17, 18]. AKT also inhibits glycogen synthase kinase 3 (GSK3) which phosphorylates PDX1 and MAFA and causes their proteasomal degradation [19-21]. Nuclear translocation of FOXO1 increases transcription of MAFA [22]. Thus, FOXO1 has opposing effects on the two transcription factors responsible for insulin transcription.

Oxidative stress and ER stress activate the downstream proteins JNK and p38MAPK [23, 24] which are responsible for downregulating insulin gene expression. JNK causes serine phosphorylation of the insulin signaling protein, IRS1, which inhibits its ability to phosphorylate AKT [25]. As a result, AKT cannot have its inhibitory actions on FOXO1 and GSK3. JNK and p38MAPK also directly phosphorylate FOXO1 and increase its nuclear translocation [18]. Oxidative stress causes translocation of MAFA from the nucleus to the cytosol. Thus, in the presence of prolonged hyperglycemia, oxidative stress and ER stress mediate the nuclear exclusion and degradation of the insulin gene transcription factors, PDX1 and MAFA, mainly through the stress-activated kinases, JNK and p38MAPK, and hence inhibit insulin gene expression.

Since AKT is responsible for controlling the activation of the important repressors of the insulin transcription factors, we only consider the effect of insulin on AKT without considering the intermediate proteins.

All the regulating actions among the proteins described in the previous paragraphs are modelled using simple mass action kinetics.

AKT is phosphorylated and activated by insulin and inhibited by JNK. The rate equations for the phosphorylation of AKT by insulin, $\mathrm{v}_{\mathrm{AKT-insulin}}$, and dephosphorylation of AKT, $v_{\mathrm{AKTp}}$, are given by:

$$v_{\mathrm{AKT-insulin}} = k_{\mathrm{AKT}} \times [AKT] \times \frac{[INS]_{\mathrm{e.c.}}}{[INS]_{\mathrm{e.c.}_{\mathrm{t=0}}}} \quad (S34)$$

$$v_{\mathrm{AKTp}} = k_{\mathrm{AKTp}} \times \left[AKT_p\right] \quad (S35)$$

where, $[AKT]$ and $[AKT_p]$ denote the concentrations of unphosphorylated and phosphorylated AKT respectively, $[INS]_{\mathrm{e.c.}}$ denotes the extra-cellular insulin concentration, $k_{\mathrm{AKT}}$ is the phosphorylation rate of AKT by insulin and $k_{\mathrm{AKTp}}$ is the dephosphorylation rate of AKT. The rate equation for the inactivation of AKT by JNK is given by:

$$v_{\mathrm{AKT-JNK}} = k_{\mathrm{AKT-JNK}} \times \left[AKT_{\mathrm{p}}\right] \times \left[JNK_{\mathrm{p}}\right] \quad (S36)$$

where, $[JNK_{\mathrm{p}}]$ denotes the concentration of phosphorylated JNK, $k_{\mathrm{AKT-JNK}}$ is the rate of inactivation of AKT by JNK.

AKT phosphorylates GSK3 and keeps it inactive. The rate equations for the phosphorylation of GSK3 by AKT, $v_{\mathrm{GSK3-AKT}}$, and dephosphorylation of GSK3, $v_{\mathrm{GSK3p}}$, are given by:

$$v_{\mathrm{GSK3-AKT}} = k_{\mathrm{GSK3}} \times [GSK3] \times [AKT_{\mathrm{p}}] \tag{S37}$$

$$v_{\mathrm{GSK3p}} = k_{\mathrm{GSK3p}} \times [GSK3_{\mathrm{p}}] \tag{S38}$$

where, $[GSK3]$ and $[GSK3_{\mathrm{p}}]$ denote the concentrations of unphosphorylated and phosphorylated GSK3 respectively, $k_{\mathrm{GSK3}}$ is the phosphorylation rate of GSK3 by AKT and $k_{GSK3p}$ is the dephosphorylation rate of GSK3.

The stress-activated kinases, JNK and p38MAPK, are activated by $H_2O_2$ ($[R_{\mathrm{h}}]$) and UPR ($U_{\mathrm{PR}}$). The rates of phosphorylation of JNK by ROS, $v_{\mathrm{JNK-ROS}}$, and UPR, $v_{\mathrm{JNK-UPR}}$, and dephosphorylation of JNK, $v_{\mathrm{JNKp}}$, are given by:

$$v_{\mathrm{JNK-ROS}} = k_{\mathrm{JNK-ROS}} \times [JNK] \times \frac{[R_{\mathrm{h}}]}{[R_{\mathrm{h}}]_{\mathrm{t=0}}} \tag{S39}$$

$$v_{\mathrm{JNK-UPR}} = k_{\mathrm{JNK-UPR}} \times [JNK] \times v_{\mathrm{UPR}} \tag{S40}$$

$$v_{\mathrm{JNKp}} = k_{\mathrm{MAPKp}} \times [JNK_{\mathrm{p}}] \tag{S41}$$

where, $[JNK]$ denotes the concentration of unphosphorylated JNK, $k_{\mathrm{JNK-ROS}}$ is the phosphorylation rate of JNK by $H_2O_2$, $k_{\mathrm{JNK-UPR}}$ is the phosphorylation rate of JNK by UPR, and $k_{\mathrm{MAPKp}}$ is the dephosphorylation rate of JNK.

The rates of phosphorylation of p38MAPK by ROS, $v_{\mathrm{p38-ROS}}$, and UPR, $v_{\mathrm{p38-UPR}}$, and dephosphorylation of p38 MAPK, $v_{\mathrm{p38p}}$, are given by:

$$v_{\mathrm{p38-ROS}} = k_{\mathrm{p38-ROS}} \times [p38MAPK] \times \frac{[R_{\mathrm{h}}]}{[R_{\mathrm{h}}]_{\mathrm{t=0}}} \tag{S42}$$

$$v_{\mathrm{p38-UPR}} = k_{\mathrm{p38-UPR}} \times [p38MAPK] \times v_{\mathrm{UPR}} \tag{S43}$$

$$v_{\mathrm{p38p}} = k_{\mathrm{MAPKp}} \times [p38MAPK_{\mathrm{p}}] \tag{S44}$$

where, where $[p38MAPK]$ and $[p38MAPK_{\mathrm{p}}]$ denote the concentrations of unphosphorylated and phosphorylated p38MAPK respectively, $k_{\mathrm{p38-ROS}}$ is the phosphorylation rate of p38MAPK by $H_2O_2$, $k_{\mathrm{p38-UPR}}$ is the phosphorylation rate of p38MAPK by UPR, and $k_{\mathrm{MAPKp}}$ is the dephosphorylation rate of p38MAPK.

AKT phosphorylates FOXO1 and keeps it in the cytosol. JNK and p38MAPK phosphorylate FOXO1 and facilitate its nuclear translocation. The rate of phosphorylation of FOXO1 by AKT, $v_{\mathrm{FOXO1-AKT}}$, is given by:

$$v_{\mathrm{FOXO1-AKT}} = k_{\mathrm{FOXO1-AKTp}} \times [FOXO1] \times [AKT_{\mathrm{p}}] \qquad (S45)$$

where, [FOXO1] denotes the concentration of unphosphorylated FOXO1 and $k_{\mathrm{FOXO1}}$ is the phosphorylation rate of FOXO1 by AKT. The rate of phosphorylation FOXO1 by JNK and p38MAPK, $v_{\mathrm{FOXO1-MAPK}}$, is given by:

$$v_{\mathrm{FOXO1-MAPK}} = k_{\mathrm{FOXO1p-MAPKp}} \times [FOXO1_{\mathrm{p}}] \times ([JNK_{\mathrm{p}}] + [p38MAPK_{\mathrm{p}}]) \qquad (S46)$$

where, $[FOXO1_{\mathrm{p}}]$ denotes the concentration of phosphorylated FOXO1, $k_{\mathrm{FOXO1p-MAPKp}}$ is the phosphorylation rate of FOXO1 by JNK and p38MAPK.

PDX1 production is increased by transcription factor MAFA and decreased by transcription factor FOXO1. FOXO1 also causes nuclear exclusion of PDX1. GSK3 causes proteasomal degradation of PDX1. The rate of production of PDX1 due to the binding of the activator transcription factor, MAFA, and repressor transcription factor, FOXO1, is given by:

$$v_{\mathrm{PDX1-prod}} = V_{\mathrm{mPDX1}} \times \frac{[MAFA]}{K_{\mathrm{MAFA-PDX1}} + [MAFA]} \times \frac{K_{\mathrm{FOXO1-PDX1}}}{K_{\mathrm{FOXO1-PDX1}} + [FOXO1]} \qquad (S47)$$

where, $[MAFA]$ is the concentration of MAFA, $V_{\mathrm{mPDX1}}$ is the maximum rate of PDX1 mRNA production, $K_{\mathrm{MAFA-PDX1}}$ is the dissociation constant of MAFA from the binding site of PDX1 promoter, and $K_{\mathrm{FOXO1-PDX1}}$ is the dissociation constant of FOXO1 from the binding site of PDX1 promoter. The rate of efflux of PDX1 from the nucleus by FOXO1 is given by:

$$v_{\mathrm{PDX1-efllux-FOXO1}} = k_{\mathrm{PDX1-FOXO1-efflux}} \times [PDX1] \times [FOXO1] \qquad (S48)$$

and the rate of proteasomal degradation of PDX1 by GSK3 is given by:

$$v_{\mathrm{PDX1-pdeg-GSK3}} = k_{\mathrm{GSK3-PDX1-pdeg}} \times [PDX1] \times [GSK3] \qquad (S49)$$

where, $[PDX1]$ is the concentration of PDX1, $k_{\mathrm{PDX1-FOXO1-efflux}}$ is the efflux rate of PDX1 from nucleus by FOXO1 and $k_{\mathrm{GSK3-PDX1-pdeg}}$ is the rate of PDX1 protein degradation by GSK3.

MAFA production is increased by transcription factors PDX1 and FOXO1. ROS causes nuclear exclusion of MAFA and GSK3 causes proteasomal degradation of MAFA. The rate of production of MAFA due to the binding of the activator transcription factors, PDX1 and FOXO1, is given by:

$$v_{\mathrm{MAFA-prod}} = V_{\mathrm{mMAFA}} \times \frac{[PDX1]}{K_{\mathrm{PDX1-MAFA}} + [PDX1]} \times \frac{[FOXO1]}{K_{\mathrm{FOXO1-MAFA}} + [FOXO1]} \qquad (S50)$$

where, $V_{\mathrm{mMAFA}}$ is the maximum rate of MAFA mRNA production, $K_{\mathrm{PDX1-MAFA}}$ is the dissociation constant of PDX1 from the binding site of MAFA promoter, and $K_{\mathrm{FOXO1-MAFA}}$ is the dissociation constant of FOXO1 from the binding site of MAFA promoter. The rate of efflux of MAFA from the nucleus by $H_2O_2$ ($[R_{\mathrm{h}}]$) is given by:

$$v_{\mathrm{MAFA-efflux-ROS}} = k_{\mathrm{ROS-MAFA}} \times [MAFA] \times \frac{[R_{\mathrm{h}}]}{[R_{\mathrm{h}}]_{\mathrm{t=0}}} \qquad (S51)$$

and the rate of proteasomal degradation of MAFA by GSK3 is given by:

$$v_{\mathrm{MAFA-pdeg-GSK3}} = k_{\mathrm{GSK3-MAFA-pdeg}} \times [MAFA] \times [GSK3] \qquad (S52)$$

where, $k_{\mathrm{ROS-MAFA}}$ is the efflux rate of MAFA from the nucleus by ROS and $k_{\mathrm{GSK3-MAFA-pdeg}}$ is the rate of MAFA protein degradation by GSK3.

Insulin mRNA production is increased by its two major transcription factors, PDX1 and MAFA. The rate of production of insulin mRNA due to the binding of the activator transcription factors, PDX1 and MAFA, is given by:

$$v_{\mathrm{INS-prod}} = V_{\mathrm{mINS}} \times \frac{[PDX1]}{K_{\mathrm{PDX1-INS}} + [PDX1]} \times \frac{[MAFA]}{K_{\mathrm{MAFA-INS}} + [MAFA]} \qquad (S53)$$

and the rate of degradation of insulin mRNA is given by:

$$v_{\mathrm{INS-deg}} = k_{\mathrm{dm-Ins}} \times [INS_{\mathrm{mRNA}}] \qquad (S54)$$

where, $[INS_{\mathrm{mRNA}}]$ is the concentration of insulin mRNA, $V_{\mathrm{mINS}}$ is the maximum rate of insulin mRNA production, $K_{\mathrm{PDX1-INS}}$ is the dissociation constant of PDX1 from the binding site of insulin promoter, $K_{\mathrm{MAFA-INS}}$ is the dissociation constant of MAFA from the binding site of insulin promoter, and $k_{\mathrm{dm-Ins}}$ is the rate of insulin mRNA degradation.

**1.7 Protein folding and unfolded protein response**

In order to restore the protein folding capacity, the unfolded protein response (UPR) is initiated. The UPR acts by decreasing translation, restoring protein folding, and causing ER-associated degradation (ERAD) of misfolded proteins [26]. In our model, we consider (i) the attenuation of protein translation by UPR, (ii) the activation of ER chaperones by UPR, and (iii) protein folding and degradation mediated by the ER chaperones.

The rate of translation of insulin mRNA to proinsulin is described by the following equation:

$$v_{\mathrm{proinsulin-prod}} = k_{\mathrm{tr-INS}} \times [INS_{\mathrm{mRNA}}] \times \frac{1}{1 + v_{\mathrm{UPR}}} \qquad (S55)$$

where, $k_{\mathrm{tr-INS}}$ is the rate of translation of insulin mRNA and $v_{\mathrm{UPR}}$ represents the UPR. This equation takes into account the attenuation of the translation process by the UPR. The rate of protein folding and degradation processes mediated by the ER chaperones are described by the rate equation taken from Graham [13]:

$$v_{\mathrm{folding-deg-chap}} = d_{\mathrm{uu}} \times [CP] \times \frac{[INS_{\mathrm{UF}}]}{[INS_{\mathrm{UF}}] + K_{\mathrm{uu}}} \qquad (S56)$$

where, $[INS_{\mathrm{UF}}]$ denotes the concentration of proinsulin (unfolded insulin), $[CP]$ is the concentration of ER chaperones, $d_{\mathrm{uu}}$ is the rate of proinsulin folding/degradation and $K_{\mathrm{uu}}$ is the proinsulin-chaperone Michaelis-Menten constant (folding/degradation).

The rates of activation of ER chaperones by the UPR, $v_{\mathrm{chap-act}}$, and the degradation of ER chaperones, $v_{\mathrm{chap-deg}}$, are described as:

$$v_{\text{chap-act}} = \frac{v_{\text{UPR}}}{1 + v_{\text{UPR}}} \tag{S57}$$

$$v_{\text{chap-deg}} = d_{\text{CP}} \times [CP] \tag{S58}$$

where, $d_{\text{CP}}$ is the rate of chaperone decay.

The rates of folding of proinsulin molecules by ER chaperones ($v_{\text{insulin-folding}}$), insulin secretion ($v_{\text{insulin-secretion}}$) and clearance of extra-cellular insulin ($v_{\text{insulin-clearance}}$) are described as:

$$v_{\text{insulin-folding}} = q_{\text{f}} \times v_{\text{folding-deg-chap}} \tag{S59}$$

$$v_{\text{insulin-secretion}} = k_{\text{IS}} \times [INS]_{\text{i.c.}} \times \frac{max\left(0, \left([\text{Ca}^{2+}]_{\text{C}} - [\text{Ca}^{2+}]_{\text{C,min}}\right)\right)}{[\text{Ca}^{2+}]_{\text{C,min}}} \tag{S60}$$

$$v_{\text{insulin-clearance}} = k_{\text{IC}} \times [INS]_{\text{e.c.}} \tag{S61}$$

where, $[INS]_{\text{i.c.}}$ is the concentration of folded insulin, $q_{\text{f}}$ is the proinsulin folding fraction, $k_{\text{IS}}$ is the rate of insulin secretion, $[\text{Ca}^{2+}]_{\text{C,min}}$ is the threshold $Ca^{2+}$ concentration necessary for insulin secretion, and $k_{\text{IC}}$ is the rate of insulin clearance. After uptake of glucose via GLUT1, the glucose is metabolized to produce ATP. The rise in ATP leads to closing of plasma membrane located ATP-sensitive $K^{+}$ (K-ATP) channels causing depolarization [2]. This facilitates the entry of $Ca^{2+}$ into the cytosol [2] which ultimately leads to exocytosis of insulin-containing granules.

The UPR is activated by the accumulation of unfolded proteins and is inhibited by the ER chaperones and it is represented as [13]:

$$v_{\text{UPR}} = \frac{\frac{[INS_{\text{UF}}]}{K_1}}{1 + \frac{[INS_{\text{UF}}]}{K_1} + \frac{[CP]}{K_2\left(1 + \frac{[INS_{\text{UF}}]}{K_3}\right)}} \tag{S62}$$

where, $K_1$ is the PERK-proinsulin dissociation constant, $K_2$ is the PERK-chaperone dissociation constant, and $K_3$ is the proinsulin-chaperone dissociation constant.
